# 3D MINFLUX combined with DNA-PAINT resolves the arrangement of Bassoon at active zones

**DOI:** 10.1101/2025.09.03.673830

**Authors:** Florelle Domart, Evelyn Garlick, Isabelle Jansen, Maria Augusta do Rego Barros Fernandes Lima, Thomas Dresbach

**Author notes:** **Correspondence** Thomas Dresbach, Institute of Anatomy and Cell Biology, University Medical Center Göttingen, Georg-August University of Göttingen, 37075 Göttingen, Germany.

## Abstract

Neurotransmitter release and membrane retrieval at active zones require precise spatial and temporal coordination relying on an intricate molecular machinery. However, the exact nano-structural organization of this machinery is not yet fully elucidated. Here, we used 3D MINFLUX combined with both spectral demixing and DNA-PAINT to analyze the positioning of the scaffolding protein Bassoon at presynaptic active zones of glutamatergic spine synapses of hippocampal neurons, achieving a localization precision of 5 nm in 3D. This approach allowed us to visualize directly the distribution of N-terminal and the C-terminal regions of Bassoon, and demonstrates that Bassoon exhibits an orientation at the active zone, where the C-terminal region is directed toward the synaptic cleft and the N-terminal region towards synaptic vesicles. Having demonstrated this spatial configuration for endogenous and recombinant Bassoon molecules, our study paves the way towards molecular-scale resolution analysis of other prominent proteins of the presynaptic release machinery.

## Introduction

Synapses are asymmetric cell-cell junctions harboring highly specialized arrangements of molecules on both sides of the synaptic cleft. These molecular machineries, comprised of synaptic scaffolding molecules and their associated binding partners, have evolved to mediate the complex events underlying neurotransmission and synaptic plasticity. This is particularly prominent on the presynaptic side: here, exocytosis of neurotransmitter from synaptic vesicles (SVs) and endocytic retrieval of SV membranes occurs in an intricately balanced manner at specialized neurotransmitter release sites, called active zones (Südhof 2012). In addition, physiological protein degradation and replenishment have to be tightly coordinated with these events (Gundelfinger et al., 2022). High levels of spatial and temporal precision are further required during assembly, maturation and remodeling of synapses, and even subtle defects may contribute to neurodevelopmental and neurodegenerative disorders (Yeo et al, 2022; Kaladiyil et al, 2025; Park et al, 2018). The way these active zone scaffolding proteins and their associated binding partners are organized at the nano-structural scale is only now beginning to yield to microscopic analysis (Tabares and Rizzoli, 2022).

The ultrastructural localization of active zone scaffolding proteins has been addressed using post-embedding immuno-gold electron microscopy (Siksou et al., 2007; Limbach et al., 2011; Holderith et al., 2012). However, labelling distinct epitopes simultaneously and obtaining 3D ultrastructural information from such dense samples is technically challenging. Super-resolution light microscopy techniques such as stimulated emission depletion (STED) microscopy and Single-Molecule Localization Microscopy (SMLM) techniques have offered an alternative approach to studying the nanoscale organization of synaptic molecules (Nosov et al., 2020). In particular, STORM (Stochastic Optical Reconstruction Microscopy) microscopy and DNA-PAINT (Point Accumulation for Imaging in Nanoscale Topography) offer sub-15 nm lateral resolution. Among these super-resolution approaches, MINFLUX (MINimal photon FLUXes) microscopy offers both lateral and axial resolution in the single-digit nanometer range (Balzarotti et al., 2017; Schmidt et al., 2021). Using this emerging technique, the positioning of active zone scaffolding proteins in photoreceptor synapses (Grabner et al., 2022), and postsynaptic scaffolding protein PSD95 in dendritic spines of primary cultured neurons (Gürth et al., 2023) have been determined with a localization precision of 5 nm. More recently, it was shown that MINFLUX can measure intramolecular distances in the sub-nanometer range in biochemically purified proteins tagged at multiple sites (Sahl et al., 2024). Thus, MINFLUX even allows for the detection of distinct regions of a given molecule. Extracting information on the distribution and intramolecular arrangement of epitopes is particularly meaningful when applied *in situ*, i.e. in the cellular environment. This task is also particularly challenging, due to the crowded environment encountered in cells, and is further complicated when the structure and arrangement of the protein investigated is not known beforehand.

Here, we employed MINFLUX microscopy to study the architecture of active zones in presynaptic nerve terminals of chemically fixed cultured hippocampal neurons, focusing on the presynaptic scaffolding molecule Bassoon, which is a particularly large active zone scaffolding molecule, comprising 3938 amino acids in the rat, and 3926 amino acids in humans (tom Dieck et al., 1998; Gundelfinger et al., 2016).

Investigating both endogenous and recombinant Bassoon, we provide a 3D analysis of the localization of an N-terminal and a C-terminal Bassoon epitope within individual active zones with a localization precision of 5 nm.

## Results

### 2 color 3D-MINFLUX imaging of synapses

Due to its nanometer scale localization precision in 3D, MINFLUX microscopy is currently the only technique capable of resolving the location of distinct epitopes of a given presynaptic protein in its cellular environment and in 3D. STORM analysis has suggested that the C-terminal region of Bassoon is located on average 30 nm closer to the postsynaptic protein Homer than its N-terminal region (Dani et al., 2010). Thus, Bassoon may be oriented with its C-terminus towards the plasma membrane and its N-terminus towards SVs.

Here, we aimed to determine the localization of these two epitopes of Bassoon directly, equipped with a 5 nm precision, in 3D using two-color MINFLUX microscopy. To achieve two-color imaging, we first used spectral demixing (Figure 1). Spectral demixing allows for the simultaneous imaging of two fluorophores with close excitation emission spectra, such as AF647 and CF680. Separation is accomplished by ratiometric imaging, where the mixed fluorescence is sent by a dichroic mirror to two detectors at wavelengths 650-685nm and 685-720 nm. Each localization is assigned to a specific fluorophore population based on the photon ratio detected by each detector (Tonnesen et al., 2011). This ratiometric method avoids chromatic aberration issues commonly found in traditional two-color SMLM (Friedl et al, 2023). To validate this approach, we triple-stained cultured neurons with phalloidin-AF488, to reveal dendritic spines, as well as a monoclonal antibody to target a more N-terminally located epitope of Bassoon comprising amino acids 738-1035 of rat Bassoon (called “N-Bassoon” in Figure 1b) and a polyclonal antibody against the postsynaptic protein Homer1 (Figure 1). Secondary antibodies were conjugated with the dyes Flux640 (for Homer1) and Flux680 (for Bassoon).

**Figure 1:**
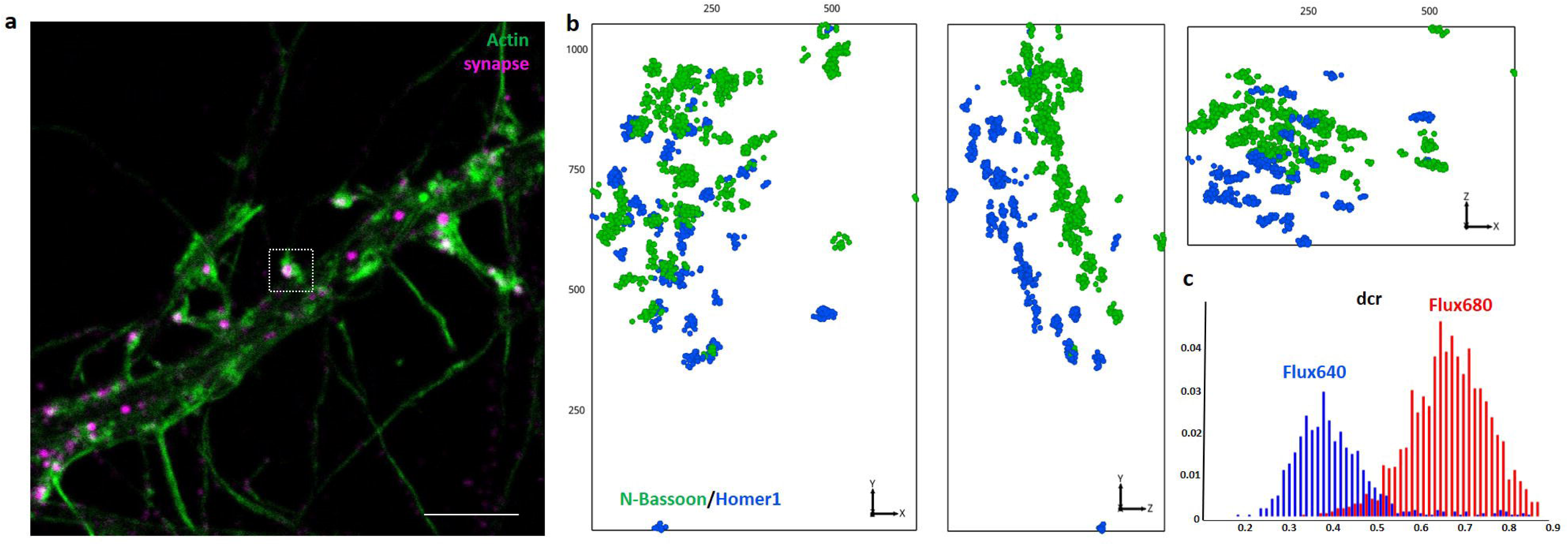
Resolution of the pre and postsynapse by 3D-Spectral demixing Minflux microscopy. a) Confocal microscopy image of a dendrite of a primary rat hippocampal neuron (DIV15) stained for actin (phalloidin-AF488, green), N terminal epitope of Bassoon (Flux680) and Homer1 (Flux640) before spectral demixing (magenta, termed synapse in the panel). Scale bar = 5 µm. b) 3D Minflux of Homer1 (blue) + Bassoon (green) in 3 different orientations after spectral demixing. The first orientation (xy) corresponds to the same orientation shown in panel a). Scales indicated along the side of the boxes in nm. c) distributions of detector channel ratio (dcr) values for Flux680 and Flux640. The locations are assigned to Flux640 or Flux680 based on the ratio of photons received by each detector.

Throughout our workflow, we imaged synapses first in confocal mode, and selected them based on both actin staining (revealing dendritic spines) and the combined emission of the signals coming from Homer and Bassoon (Figure 1a), ensuring that all synapses we imaged indeed were excitatory spine synapses. We then recorded MINFLUX data from these synapses. We explored different pairs of fluorophores and identified the pair Flux680 and Flux640 as optimal in terms of blinking kinetics, offering on average a localization precision in 3D of 5.7 ± 0.1 nm (mean ± SEM) for Flux680 and 5.6 ± 0.1 nm (mean ± SEM) for Flux640, as evidenced by the reconstructed images for Bassoon and Homer1 (Figure 1b, Figure S1). In addition, the histogram of detector channel ratio (dcr) confirmed adequate spectral separation of the two fluorophores (Figure 1c). Note that in the 3D-MINFLUX images in the left panel of Figure 1b, labelled “xy” in the panel, shows the orientation of the synapse as in the confocal view, while the middle and the left panels show 90-degree flips (yz and xz) relative to the original orientation. In this example, the middle panel (yz) shows a side view of the synapse with the synaptic cleft. The effectiveness of the ratiometric method based on a clear-cut fluorophore assignment is particularly evident in this side view of the synapse (yz orientation) (Figure 1b).

### 3D-MINFLUX spectral demixing resolves the orientation of Bassoon at the active zone

After validating imaging of pre- and postsynaptic proteins with a 5 nm localization precision and effective spectral demixing, we co-labelled cultured neurons with the monoclonal antibody herein called “N-Bassoon” and a polyclonal antibody raised against the C-terminus of Bassoon, called “Bassoon-C” here. As in Figure 1, we selected synapses by confocal microscopy. Figures 2a shows actin staining and the combined signal from Flux640 for Bassoon-C and Flux680 for N-Bassoon. Figure 2b displays 3D-MINFLUX images of the three orientations (xy, yz, and xz), starting with the original orientation of the confocal image (xy). In the “yz” orientation (Figure 2b), a spatial segregation is observed between the accumulation sites of the Flux640 signal in purple and the Flux680 signal in green (Figure 2b). Figure 2c displays the histogram of dcr with the fluorophore assignment.

**Figure 2:**
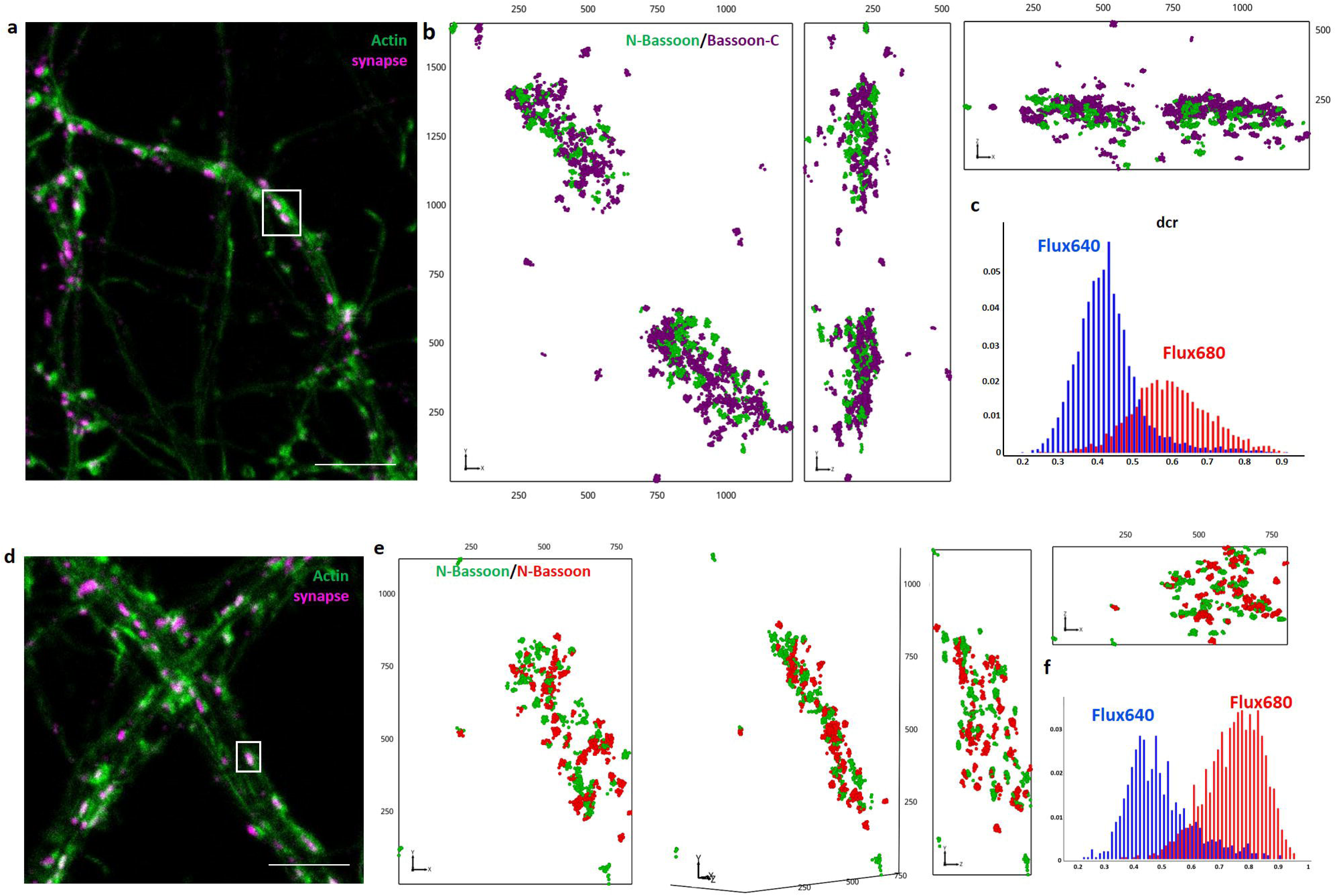
Organisation of N-Bassoon and Bassoon-C shown by 3D-Spectral demixing Minflux microscopy. a) Confocal microscopy image of a dendrite of primary rat hippocampal neurons DIV15 stained for actin (phalloidin-AF488, green), N-terminal epitope of Bassoon (N-Bassoon, Flux680) and C-terminal epitope of Bassoon (Bassoon-C, Flux640) before spectral demixing (magenta, termed synapse in the panel). Scale bar = 5 µm. b) 3D Minflux of Bassoon-C (purple) + N-Bassoon (green) in 3 different orientations after spectral demixing. Scales indicated along the side of the boxes in nm. c) distribution of dcr values for Flux680 (N-Bassoon) and Flux640 (Bassoon-C). The locations are assigned to Flux640 or Flux680 based on the ratio of photons received by each detector. d) Confocal microscopy image of a dendrite of primary rat hippocampal neurons DIV15 stained for actin (phalloidin-AF488, green), and 2 primary antibodies directed against the N-terminal epitope of Bassoon (Flux680 and Flux640) before spectral demixing (magenta, termed synapse in the panel). Scale bar = 5 µm. e) 3D Minflux of N-Bassoon (green) + N-Bassoon (red) in 4 different orientations after spectral demixing. Scales indicated along the side of the boxes in nm. f) distribution of dcr values for Flux680 (N-Bassoon) and Flux640 (N-Bassoon).

To confirm whether these observations faithfully reflect the nano-organization of Bassoon, we double-stained the N-terminal region of Bassoon using two distinct primary monoclonal antibodies. We acquired 3D MINFLUX images and confirmed effective spectral separation (Figure 2d,e,f). Unlike the staining for N-Bassoon and Bassoon-C, we observed a rather random orientation between the two distinct N-Bassoon antibodies (Figure 2e), as expected for two antibodies binding to the same epitope. Thus, our observations confirm the previous findings obtained from STORM analysis suggesting a distinct orientation of Bassoon at active zones (Dani et al, 2010) and additionally provide a direct control through the double staining of the same epitope.

To facilitate the visualization of the data shown in Figures 1 and 2 we used only the centroid of each burst of localizations, performed 3D analysis and selected the top view and side view of the synapse (Figure 3). Bassoon and Homer1 serve as scaffolding proteins located at the active zone and postsynaptic density, respectively. Thus, by orienting the reconstructed 3D image of N-Bassoon and Homer1 signals in a right way, the synaptic cleft can be revealed (Figure 3a). Indeed, rotating the 3D reconstructed images showed that the image surface covered by Homer1 or Bassoon signals was thinnest when the synaptic cleft was rotated into the viewing plane, corresponding to a side view of the synapse. Subsequently, images acquired for the N-Bassoon and Bassoon-C (Figure 3b), and control images with the two different antibodies raised against N-Bassoon (Figure 3c), were oriented to achieve the thinnest arrangements, indicative of a side view orientation. The side view aspect of N-Bassoon and Bassoon-C confirmed the spatial separation between the two epitopes. In contrast, side view images of doubly labelled N-Bassoon showed again a non-uniform orientation. This confirms that the spatial separation observed between N-Bassoon and Bassoon-C is not a technical artifact, but instead accurately reflects the nano-organization of Bassoon at the active zone.

**Figure 3:**
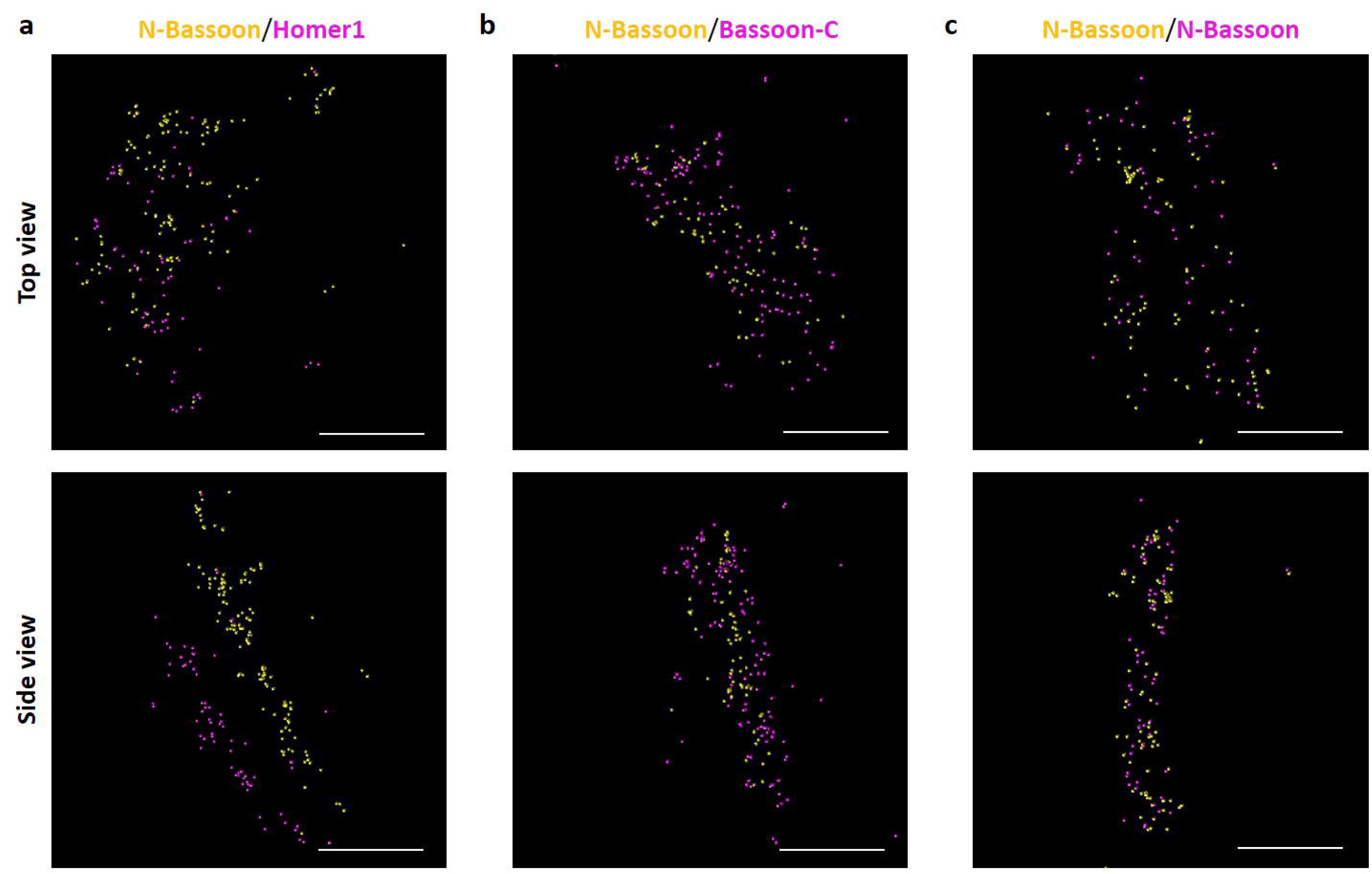
Rendering of 3D-Minflux of Bassoon and Homer data after extraction of the centroid of each trace and showing the top view and side view synapses from figures 1-3. a) N-Bassoon and Homer1. b) N-Bassoon and Bassoon-C. c) N-Bassoon and N-Bassoon. Scale bars = 200 nm. Puncta size = 7 nm.

### 3D-DNA-PAINT complements Spectral Demixing

Spectral demixing is based on the simultaneous labelling of epitopes with distinct fluorophores. An inherent technical limitation of this approach, in particular in a crowded environment, could be energy transfer from one fluorophore to another. Fluorophores that could, in principle, be localized using MINFLUX may remain undetectable due to energy transfer interactions with adjacent fluorophores situated within less than 10 nm (Helmerich and al, 2022). To rule out such a possibility, we investigated the localization of Bassoon epitopes with an additional method that relies on sequential imaging of two epitopes labelled with the same fluorophore, i.e. Exchange-PAINT. DNA-PAINT involves fluorophores coupled to short DNA strands (imagers) that transiently bind to complementary DNA strands (docking strands) attached to antibodies detecting an epitope in a target protein, and has previously been demonstrated in MINFLUX microscopy (Ostersehlt et al, 2022). The transient binding of imagers mimics the stochastic blinking effects used, e.g., in STORM microscopy. After recording signals from the first epitope samples are washed to remove the first imager and a second imager is applied, followed by recording signals from the second epitope. This strategy makes it possible to detect two targets sequentially with the same fluorophores, thus avoiding both chromatic aberrations and fluorophore interactions. In addition, to achieve closer proximity of fluorophores to their targets, we utilized docking strands conjugated to nanobodies. Using this technique, we obtained a precision of localization below 5.5 nm in 3D similar to the precision of localization obtained with spectral demixing (Figure S2).

As this approach relies on a washing step before application of the second imager, it can induce sample drift between the acquisitions. Here, we implement a method using the MINFLUX systems beamline monitoring function to correct for these shifts (see Material and Methods). We used immobile gold beads present in the sample for re-registration of the field of view after washing away the first imager and applying the second. To validate this method, we employed it on Nuclear Pore Complexes (NPC), which are well-characterized biological structures commonly used for calibration of SMLM measurements. We used a U2OS cell line stably expressing the nucleoporin NUP96 tagged with mEGFP (Thevathasan et al., 2019). We detected the GFP using an anti-GFP nanobody conjugated to a DNA-PAINT docking strand, and an imager coupled to ATTO655. We then removed the imager by washing the sample on the microscope stage, applied another batch of the same imager, and recorded again. With a z-scaling factor of 0.7 the mean absolute error of bead (MAE) was determined to be 2.95 ± 2.86 nm (mean ± SD) after drift correction, a value which is below the precision of localization and below the intra and inter molecular distances analysed in this study. Figure 4a indicates the xy and xz positions of beads before and after applying the drift correction. Figure 4b shows the xy and xz projections of the nuclear pore complex (NPC) images from the first and second acquisitions, displaying their spatial overlap before and after drift correction.

**Figure 4:**
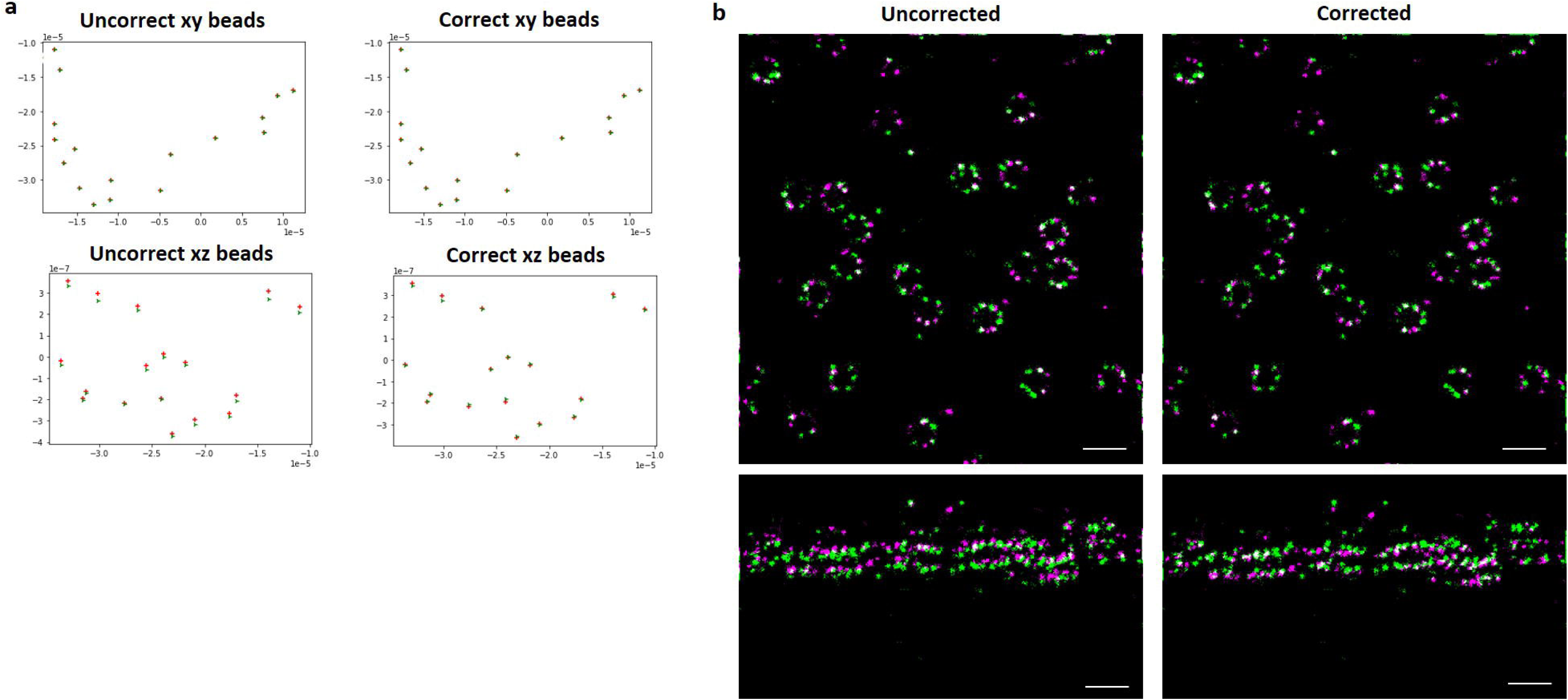
Validation of the post washing drift correction method on nucleopores (GFP tag) a) Determination of selected beads positions before and after drift correction. b) 3D-DNA-PAINT Minflux on nucleopore. After the first acquisition (shown in green) the samples were washed and the same imager was put again before a second acquisition (shown in magenta). Scale bars = 200 nm. The colocalization before and after the washing step is shown with the xy projection and xz projection before and after applying the post washing drift correction. Scale bars in m.

After drift correction, the images for the first and the second imager are overlaid with an error below the precision of localization. Thus, this error is negligible and indicates that we can perform exchange-PAINT combined with MINFLUX while fully leverage the high localization accuracy achieved by 3D-MINFLUX

### 3D-MINFLUX combined with DNA-PAINT reveals the orientation of Bassoon, with the C-terminal facing the post-synapse

We subsequently employed this method as an alternative strategy to confirm that Bassoon adopts a specific orientation within the active zone. To bring the fluorophore closer to the target, we used nanobodies, rather than secondary antibodies, directed against either mouse or rabbit primary antibodies and with specific docking sites. Synapses were identified in confocal microscopy based on actin staining and Shank2 as postsynaptic marker (Figure 5a,c). We sequentially imaged the N-terminal and the C-terminal of Bassoon, incorporating a washing step and applying post-washing drift correction to the second image (Figure 5b,d). Panels 5b display the total localizations detected in three orientations (yx, zx, and yz) for the same synapse, with Bassoon-C in purple and N-Bassoon in green, as well as the orientation where the structure is thinnest, corresponding to the side view of the synapse. The data were obtained with mean of localization precisions of σx = 5.6 nm, σy = 5.3 nm and σz = 3.7 nm for the C-terminal and σx = 5.4 nm, σy = 5.2 nm, σz = 3.4 nm for the N-terminal of Bassoon (Figure S2). In the side view orientation, we observed an accumulation of localizations in purple on one side of the active zone, corroborating the data obtained through spectral demixing. We then imaged N-Bassoon stained with two primary antibodies (Figure 5d), and the 3D MINFLUX images revealed no specific orientation but rather a random distribution aligning with what we observed in spectral unmixing data (Figure 5d). We did not observe high degree of colocalization between the two N-Bassoon images, suggesting competition between the two primary antibodies directed against the N-terminal region of Bassoon.

**Figure 5:**
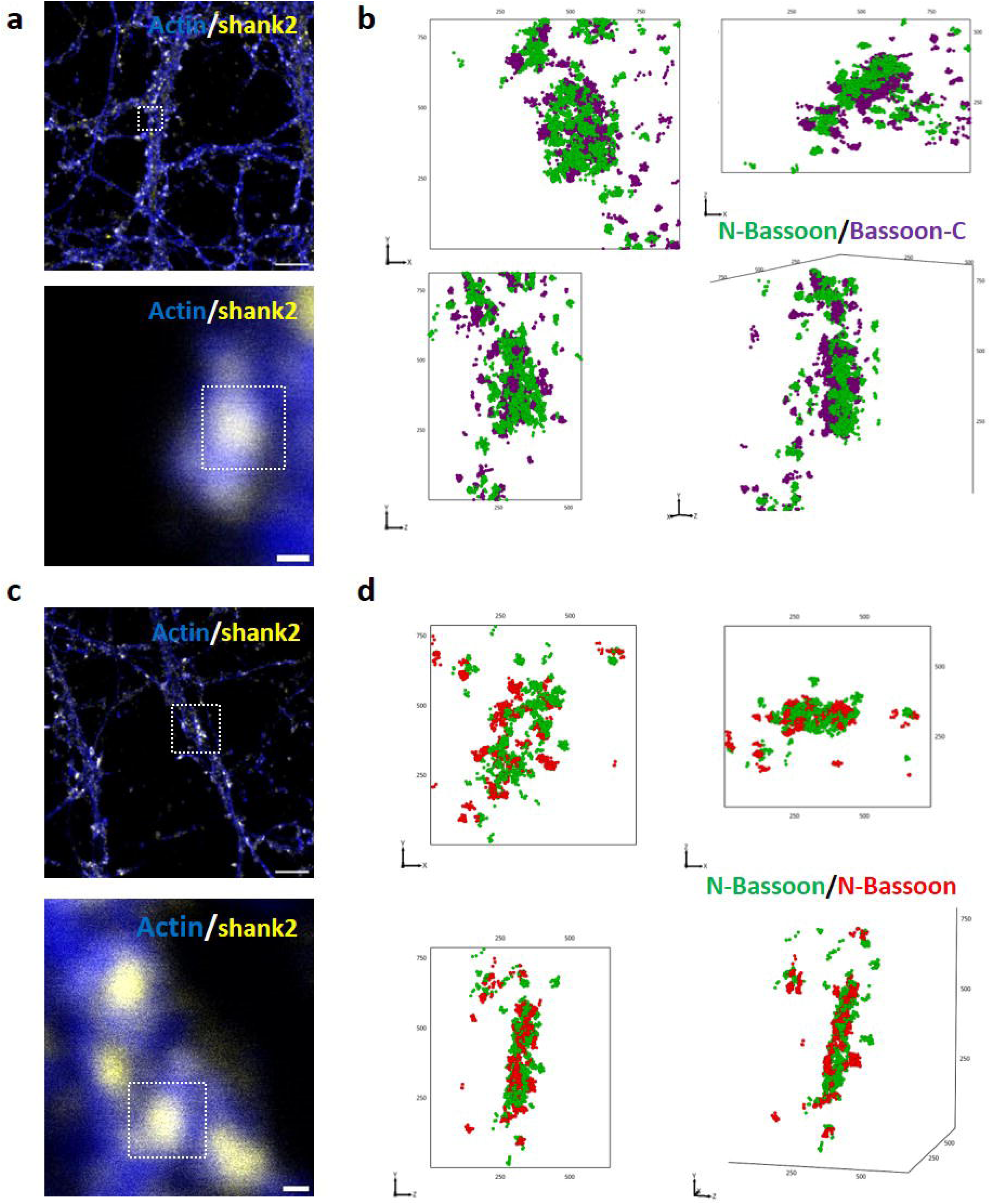
3D-DNA-Paint on endogenous highlights the orientation of Bassoon at the synapse. a) Confocal microscopy image of a dendrite of primary rat hippocampal neuron (DIV14) stained for actin (AF488, blue) and shank2 (Cy3, yellow). Scale bar = 5 µm. And confocal zoom on the synapse imaged by DNA-PAINT. Scale bar = 200 nm. b) 3D-DNA-PAINT Minflux of N-Bassoon (green) and Bassoon-C (purple) in 3 different orientations (yx, zx and yz) and in side view. Scales indicated along the side of the boxes in nm. c) confocal microscopy image and zoom of a control synapse of a primary rat hippocampal neuron (DIV14) stained for actin (AF488, blue), shank2 (Cy3, yellow). d) 3D-DNA-PAINT Minflux of N-Bassoon (green) and N-Bassoon (red) in 3 different orientations (yx, zx, yz) and in side view. Scales indicated along the side of the boxes in nm.

Extracting the centroid of each trace revealed that the separation between the C and N termini of Bassoon was more distinct using DNA-PAINT compared to the data obtained through spectral demixing (Figure 6a,b; compare to Figure 3b). Next, we aimed to determine the orientation of the termini of Bassoon with respect to the post-synapse directly using our DNA-PAINT / 3D-MINFLUX approach. To this end, we employed a rabbit anti-Homer1 antibody as a postsynaptic marker in addition to the mouse anti N-Bassoon and the rabbit anti Bassoon-C antibody. As previously, the rendered images display the centroids of each traces (Figure 6c,d). In this configuration, the anti-rabbit nanobody recognizes both the anti-Homer1 and the anti-Bassoon-C antibody. When we co-stained Bassoon and Homer1 the synaptic cleft was resolved in 3D MINFLUX images, eliminating the requirement for primary antibodies from different species to distinguish presynaptic from postsynaptic proteins. This methodological advantage enabled us to generate two-color DNA-PAINT images targeting three different molecules. We sequentially imaged N-Bassoon (yellow in Figure 6c) followed by washing and imaging of Homer 1 and Bassoon-C (purple in Figure 6c). The images were then rendered using the centroids of the burst of localizations, and drift correction was applied as before. Side view images revealed that indeed, the C-terminus of Bassoon faces the postsynapse, while the N-terminus of Bassoon is located farther away from the postsynapse (Figure 6c). As expected, this separation was not visible when the N-terminus of Bassoon was labelled with two distinct anti N-Bassoon antibodies (Figure 6d).

**Figure 6:**
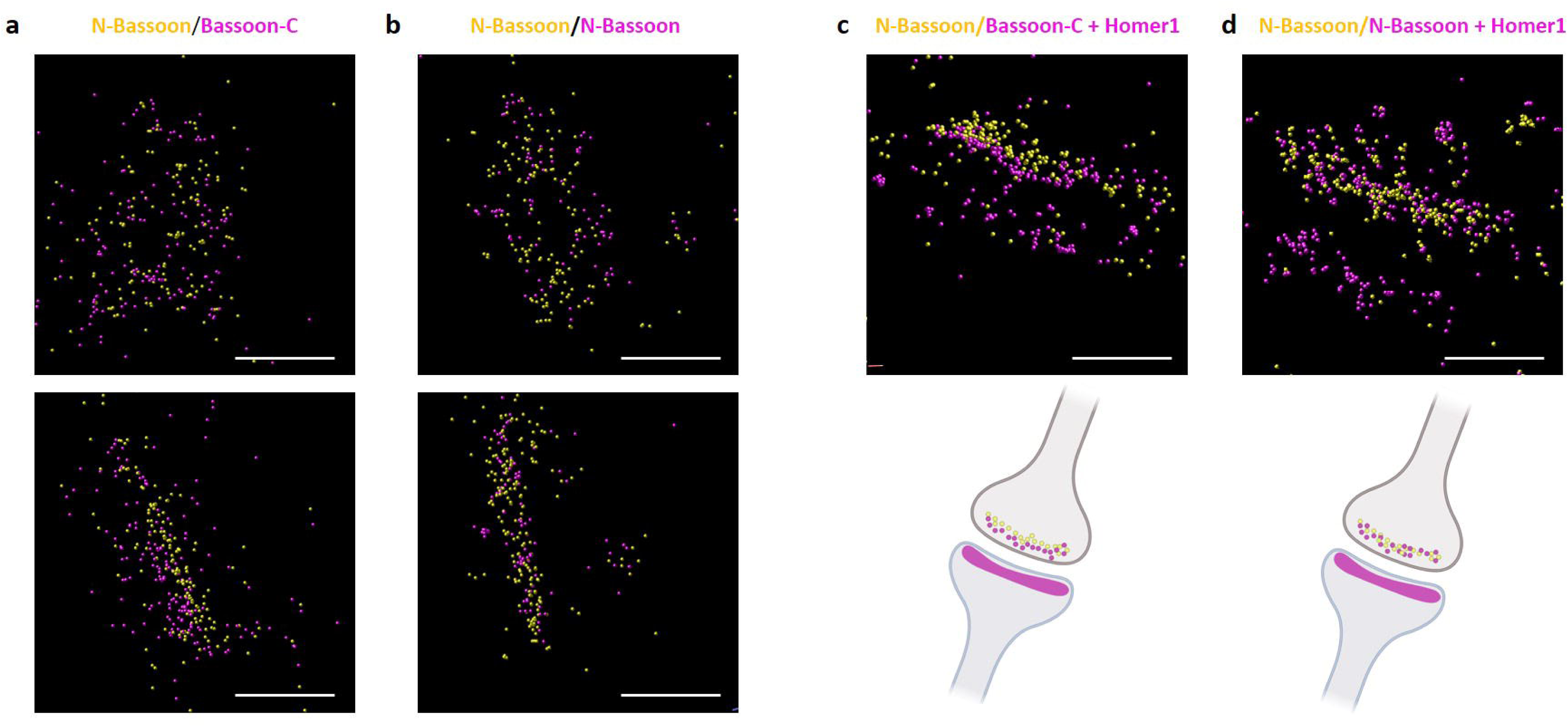
Rendering of 3D-DNA-PAINT Minflux data of endogenous Bassoon after extraction of the centroid of each traces and showing top view and side view synapses. a) Rendering of 3D-DNA-PAINT Minflux data of endogenous Bassoon, N-Bassoon and Bassoon-C. b) Rendering of 3D-DNA-PAINT Minflux data of endogenous N-Bassoon in two consecutive acquisitions and Homer1 after extraction of the centroid of each traces showing side view synapses. c) representative synapse with 2 colors N-Bassoon and Bassoon-C with the Bassoon-C facing Homer. d) representative synapse with 2 colors N-Bassoon and Homer1 accompanied by a schematic depiction illustrating the spatial distribution of N-Bassoon from two consecutive acquisitions and in relation to Homer1. Scale bars = 200 nm. Size of puncta = 7 nm.

### Organization of endogenous Bassoon at the active zone

To examine the organization of the N-terminal region of Bassoon relative to the C terminal region on a more quantitative level, we measured the volume occupied by N-Bassoon and compared it to that of Bassoon-C at 13 distinct active zones (Figure 7a). In addition, because variations in area observed in top-view synapses, despite comparable volumes, could indicate greater flexibility in one region compared to the other, we also measured the area occupied by N-Bassoon and Bassoon-C in top-view orientated synapses. We found no significant differences in either the expression volumes (Figure 7a), or the expression areas in top-view orientation (Figure 7b), suggesting a similar organization of the N-terminal and C-terminal regions of Bassoon. These results are supported by the finding that the density of localization for N-Bassoon and Bassoon-C were similar (Figure 7c). Bassoon is thought to form a dense molecular network at the active zone (AZ). However, the precise number of Bassoon molecules at the AZ has yet to be quantitatively measured on a single synapse level. Previous insights based on biochemical estimations suggested a copy number of around 450 per AZ (Wilhelm et al, 2014). Determining the number of signals at individual synapses yielded an average of 155.2 ± 16.11 Bassoon copies revealed per AZ (mean ± SEM, n = 21 AZ) (Figure 7d). It is important to emphasize that this value may be an under-estimate due to limitations in labelling efficiencies and detection efficiency. Nevertheless, thanks to the precision of our 3D-MINFLUX approach, we can report a lower bound for the copy number of Bassoon per active zone (AZ).

**Figure 7:**
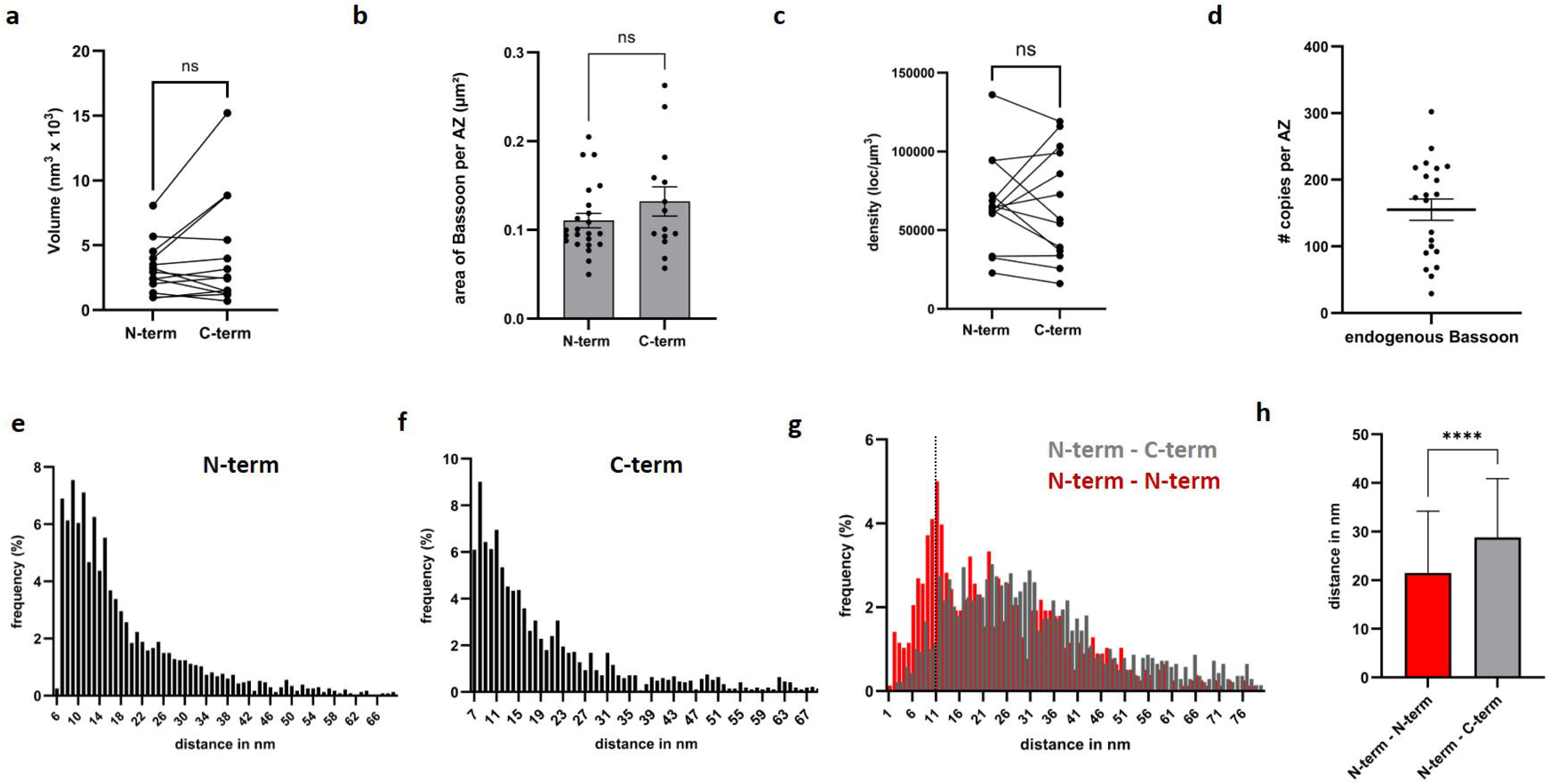
characterisation of endogenous Bassoon distribution at the Active Zone. a) Volume of expression of N-term and C-term of Bassoon correlated for each AZ (N = 13 AZ). b) Area of expression of Bassoon in synapse in top view for N-term (N = 23 AZ; 0,11 ± 0,008 µm² (mean ± SEM)) and C-term (N = 14 AZ; 0.13 ± 0.016 µm²). c) Density of localization per µm3 for N-term and C-term Bassoon and correlated for each AZ (N = 13 AZ). d) Number of Bassoon detected per AZ is about 155.2 ± 16.11 copies (Mean ± SEM). e) frequency of distribution of NND between N-term of Bassoon (n = 2334) with a median of 14.10 nm. f) frequency of distribution of NND between Basson-C (n = 2675) with a median of 14.60 nm. g) frequency of distribution of NND between N-Bassoon and Bassoon-C (grey) (n= 1387) and N-Bassoon in first acquisition compared to N-Bassoon in second acquisition (n = 780). h) distance in nm between N-Bassoon and N-Bassoon re-imaged in red (n = 780; median = 21.50; 75% quartile = 34.20; 25% quartile = 11.45) and N-Bassoon and Bassoon-C in grey (n= 1387; median = 28.80 nm; 75% quartile = 40.90; 25% quartile = 19.90; Mann Whitney and Kolmogorov-Smirnov, p-value <0,001)

Next, we determined the NND between two individual N-Bassoon molecules detected in one single acquisition (Figure 7e) and between two individual Bassoon-C detected in another single acquisition (Figure 7f). The results revealed a comparable distribution pattern for N-Bassoon and Bassoon-C. Furthermore, a Mann-Whitney test comparing the distributions of N-Bassoon (2334 individual NND) and Bassoon-C (among 2675 NND) showed no statistically significant difference (p-value = 0.37). This finding confirmed that the distribution for both sides of Bassoon is similar, validating our previous results. Additionally, the results revealed a preferred interspacing between two N-termini and between two C-termini at 10 nm, with a median distance of 14.60 nm.

Finally, we aimed to analyze the segregation of N-Bassoon and Bassoon-C observed in the 3D-MINFLUX reconstructions (see Figures 5 and 6) in a quantitative way. To this end, we performed NND analysis between N-Bassoon and Bassoon-C. We observed a median distance of 28.80 nm between N-Bassoon and Bassoon-C (Figure 7g,h). Then, we performed two sequential acquisitions of N-Bassoon and measured the NND between the two acquisitions (median = 21.70 nm). Imaging the N-terminus twice, the distribution of separation distance between N-termini from the first recording and N-termini from the second recording reveals a first peak at 9.8 ± 3.8 nm (mean ± SD). A Gaussian Mixture Model (GMM) analysis suggested additional peaks spaced at about every 11 nm (respectively at 21.2 ± 4.1 nm, 32.6 ± 4.5 nm, 45.1 ± 5.1 nm, and 62.2 ± 8.5 nm) (Figure S3). This periodicity may reflect the distance of the first, second, third and fourth closest bassoon copies and could be attributed to incomplete detection of bassoon copies between the two recordings. Importantly, these 10 nm intervals are consistent with the observed inter-protein spacing. In contrast, the distribution of separation distances between N-termini and C-termini follows a simple Gaussian profile (Figure 7g). Furthermore, the distribution of NND frequencies revealed a shift between N-Bassoon in sequential recordings versus N-Bassoon and Bassoon-C (Figure 7g). Moreover, the Mann-Whitney test as well as Kolmogorov-Smirnov tests confirmed a highly significant difference between these two distributions (Mann-Whitney, p-value < 0.0001; KS-statistic: 0.19, p-value: 8.36e-17) (Figure 7h). These results confirm that the N-terminal and C-terminal regions of Bassoon present a specific orientation at the AZ.

To better understand this change in N-Bassoon distribution across sequential recordings, we quantified the number of localizations over time in both acquisitions. Our analysis revealed a decline in locations from the first recording, with a measurable difference of up to 70% fewer locations during the first 15 minutes between the two datasets (Figure S4). This decrease may be attributed to photodamage affecting the DNA strand coupled to the nanobody, which might artifactually increases the apparent molecular distances between the first and second acquisitions when the same target is visualized twice. Additionally, we examined this distribution in sequential N-Bassoon and Bassoon-C recordings. While a reduction in localizations was observed over time during each acquisition, the second recording consistently began at the same localization level as the first. This suggests that photodamage occurs within a confined region surrounding the fluorophore, without significantly impacting the opposite side of Bassoon. These findings indicate that the measured distance between N-Bassoon and Bassoon-C is less impacted by photodamage than the control distance. However, given the limitations of labelling efficiency, the measured distance is likely an overestimate. The findings confirm that Bassoon adopts a specific orientation at the synapse; however, the actual spatial separation between the two epitopes is likely shorter than previously estimated.

### Using dually tagged recombinant Bassoon to minimize linkage error

To validate these results while minimizing the linkage error, we engineered a recombinant Bassoon construct expressing an intramolecular monomeric GFP near the N-terminus (downstream of amino acid 95) and an ALFA tag at the C-terminus, based on our previous study using dually tagged Bassoon versions (Ghelani et al.,2021). This construct was expressed in hippocampal cultures, and we verified its presynaptic localization and the absence of aberrant overexpression of Bassoon, as evidenced by the colocalization with VGluT1 and the comparable expression levels in transfected and non-transfected synapses (Figure S5). We used nanobodies for DNA-PAINT directed against the GFP and ALFA tags and performed 3D-MINFLUX (Figure 8). The data were collected with mean localization precisions of σx = 5.7 nm, σy = 5.7 nm, and σz = 3.6 nm for ALFA, and σx = 5.6 nm, σy = 5.4 nm, and σz = 3.3 nm for GFP, therefore with a localization accuracy comparable to that obtained for endogenous Bassoon. The selection of transfected synapses was based on the colocalization between the postsynaptic protein Homer1 and the GFP autofluorescence of recombinant Bassoon in confocal imaging (Figure 8a,b). Figure 8c displays three orientations (yx, yz, and zx), with yx corresponding to the orientation of the synapse in the confocal image. The yz and zx orientations distinctly show the spatial relationship between Bassoon-C (purple) and N-Bassoon (green). This observation was corroborated by rendering images using the centroids of traces (Figure 9), confirming our results obtained through indirect labeling of endogenous Bassoon. To verify that the observed orientation is not an artifact resulting from incomplete drift correction, the GFP tag was imaged a second time after a washing step. Thus, the two GFP acquisitions followed the same workflow as the acquisition sequence involving GFP followed by ALFA (Figure 9b). Unlike the shift observed between GFP and ALFA, repeated imaging of GFP did not lead to any discernible spatial shift between the two acquisitions excluding an artefact and confirming the organization of Bassoon.

**Figure 8:**
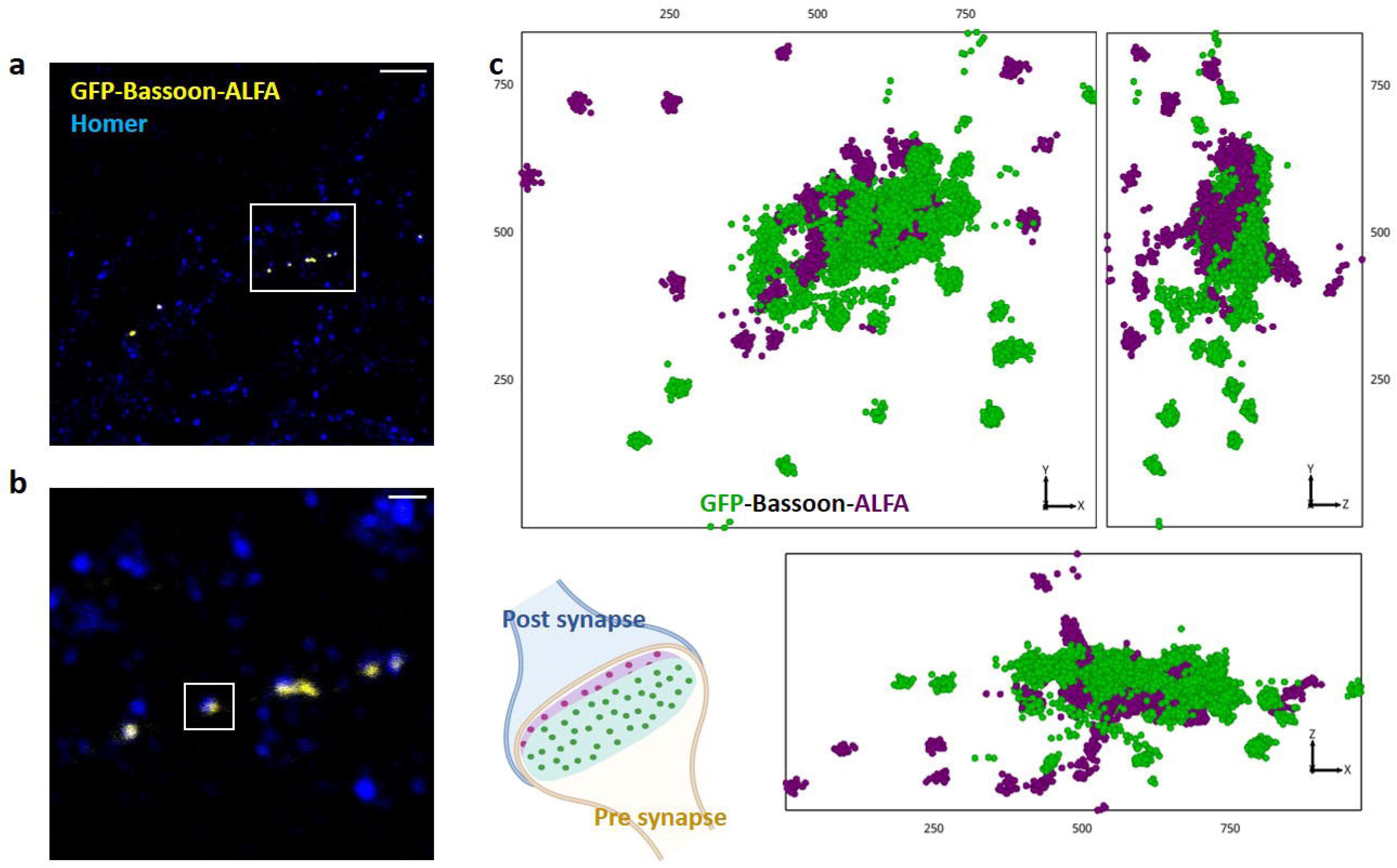
3D-DNA-Paint on GFP-Bassoon-ALFA highlights the orientation of Bassoon at the synapse. a) Confocal microscopy image of a dendrite of a primary rat hippocampal neuron (DIV14) transfected with GFP-Bassoon-ALFA and stained for Homer1 (AF488, green). Scale bar = 5 µm. b) Confocal zoom on the transfected synapse imaged by DNA-PAINT. Scale bar = 1 µm. c) 3D-DNA-Paint Minflux of GFP (green) and ALFA (purple) in 3 different orientations. Scales indicated along the side of the boxes in nm.

**Figure 9:**
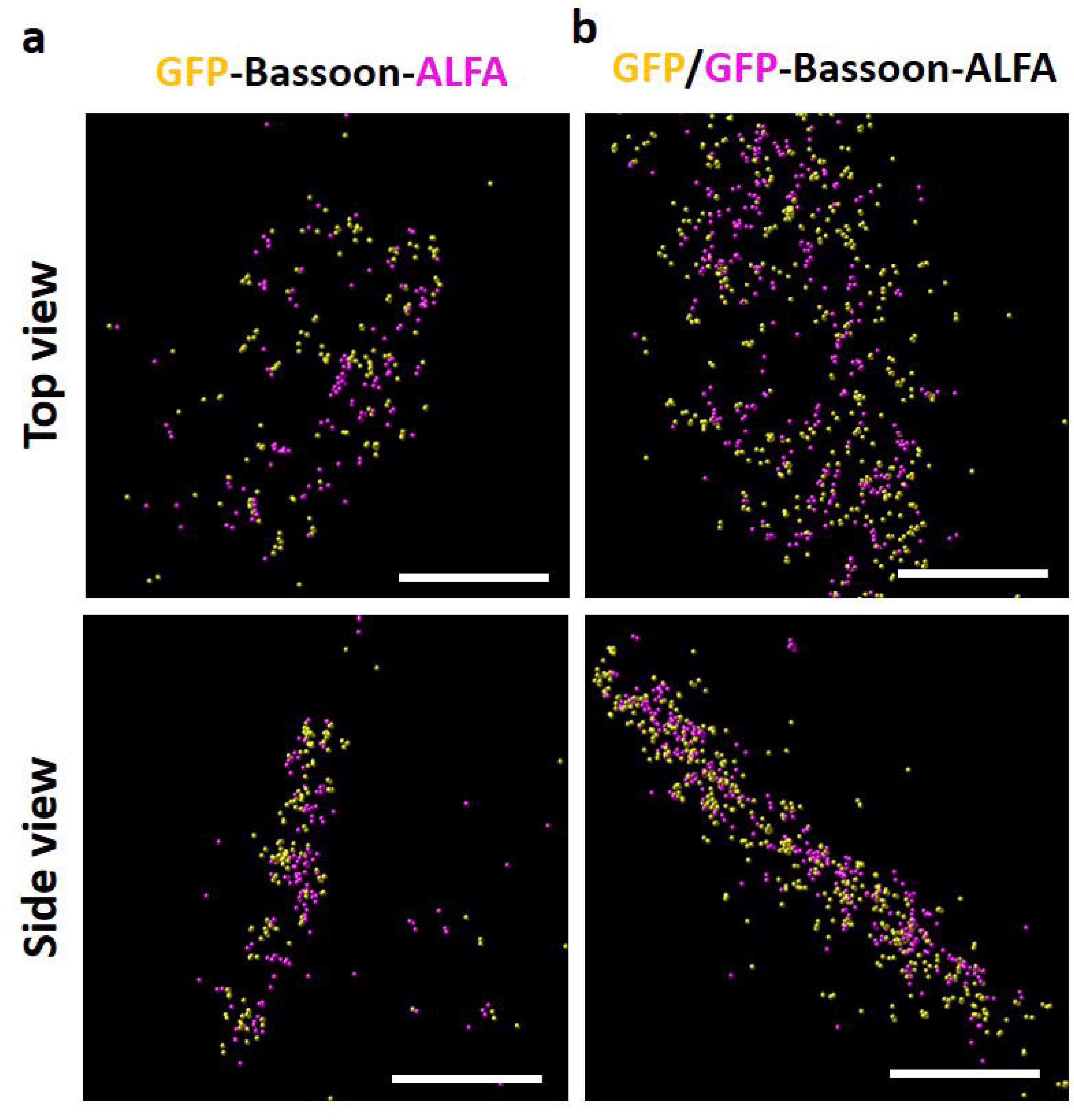
Rendering of 3D-DNA-PAINT Minflux data of recombinant Bassoon and endogenous Bassoon after extraction of the centroid of each traces and showing top view and side view synapses. a) Recombinant Bassoon GFP-Bassoon-ALFA. b) endogenous Bassoon, N-Bassoon and Bassoon-C. Scale bars = 200 nm. Size of puncta = 7 nm

### Organization of recombinant GFP-Bassoon-ALFA at the active zone

To investigate the organization of recombinant Bassoon at the active zone, we performed a volume-metric analysis of the GFP or ALFA signals using DBSCAN analysis with PoCa software (Figure 10a). We also compared the area of GFP and ALFA expression when visualizing the synapse in a top view orientation (Figure 10b) and quantified the localization density in both case (Figure 10c), as previously done for endogenous Bassoon. Our findings showed no significant differences between GFP and ALFA for these parameters, indicating a similar organization on both sides of Bassoon as found for endogenous Bassoon. Using clustering analysis based on NND applied to GFP, which is located at the N-terminal of Bassoon, we estimated the median number of recombinant Bassoon copies imaged per active zone to be 142 copies (n = 23 AZ) (Figure 10d). Additionally, as with endogenous Bassoon, we performed NND analysis between two detected GFP tags within the same acquisition, yielding a median distance of 16.10 nm (N = 15 synapses, n = 2348 NND) and two detected ALFA tags from the corresponding images, yielding a median distance of 15.10 nm (N = 15 synapses, n = 2998) (Figure 10f).

**Figure 10:**
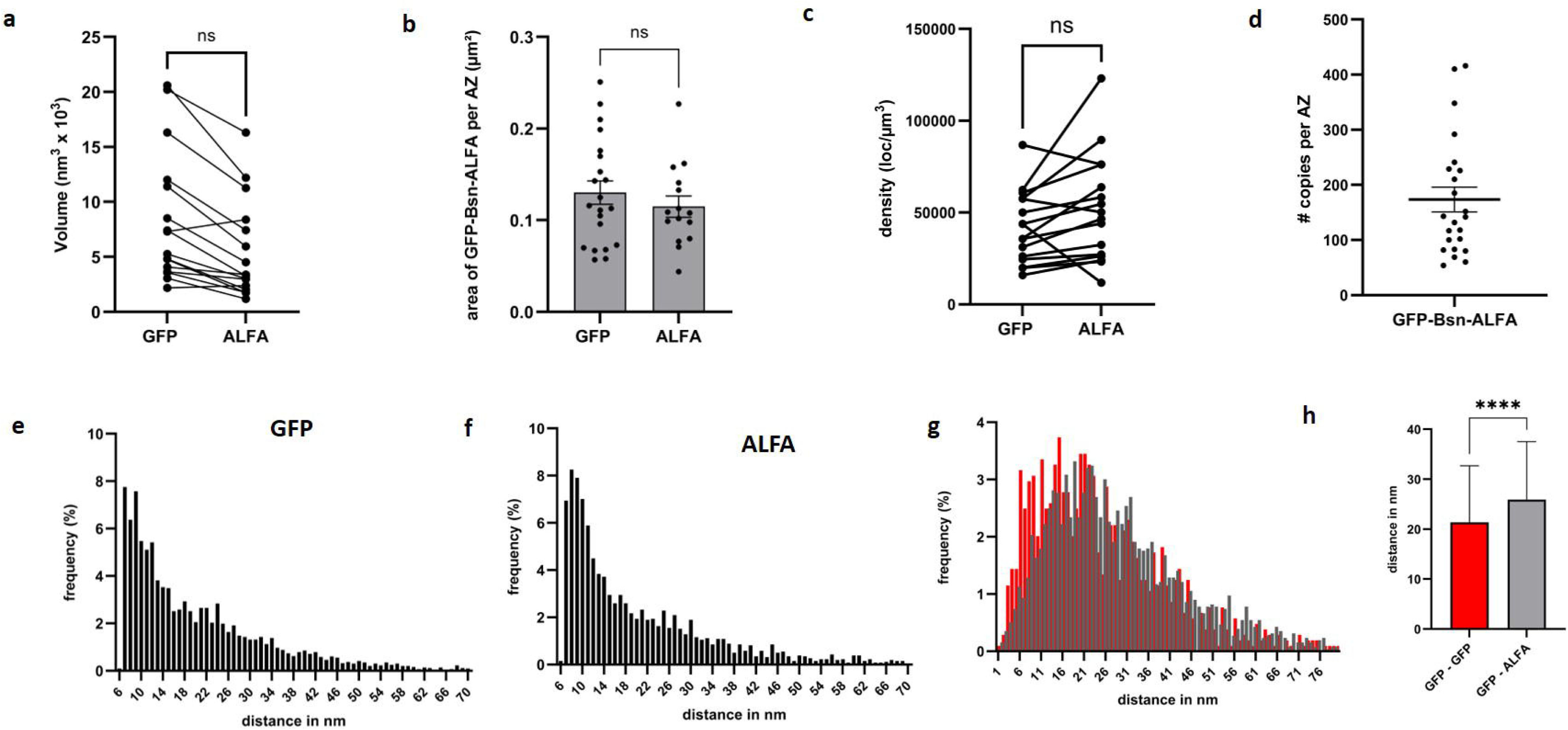
characterisation of recombinant Bassoon distribution at the Active Zone. a) Volume of expression of GFP-tag and ALFA-tag from recombinant GFP-Bassoon-ALFA correlated for each AZ (N = 16 AZ). b) Area of expression of GFP-Bassoon-ALFA in synapse in top view for GFP-tag (N = 21 AZ; 0.13 ± 0.013 µm² (mean ± SEM)) and ALFA-tag (N = 15 AZ; 0.11 ± 0.011 µm²). c) Density of localization per µm3 for GFP-tag and ALFA-tag and correlated for each AZ (N = 16 AZ). d) Median number of Bassoon detected per AZ is 142 copies. e) frequency of distribution of NND between GFP-tag (n = 4332) with a median distance of 16.10 nm. f) frequency of distribution of NND between ALFA-tag (n = 2580) with a median distance of 15.10 nm. g) frequency of distribution of NND between GFP-tag and ALFA-tag (grey) (n= 2561) and GFP-tag in first acquisition compared to GFP-tag in second acquisition (n = 1044). h) distance in nm between GFP-tag and GFP-tag re-imaged in red (n = 1044; median = 21.40; 75% quartile = 32.69; 25% quartile = 12.86) and GFP-tag and ALFA-tag in grey (n= 2561; median = 25.90 nm; 75% quartile = 37.52; 25% quartile = 17.10; Mann Whitney and Kolmogorov-Smirnov, p-value <0,0001).

The NND analysis between GFP and ALFA tags showed a median distance of 25.90 nm (N = 15 synapses, n = 2561). This is in the same range albeit slightly shorter than the distance we obtained with DNA-PAINT on endogenous Bassoon (Figure 10g,f). We then imaged the GFP tag twice within the same synapse finding a median NND of 21.40 nm (N = 9 AZ, n = 1044), similar to the distance found between two N-Bassoon molecules in first recording vs second recording. Despite the limitation in the control condition, might give us an overestimate for the distance (see above; Figure S4), the difference between distribution and distances found between GFP-ALFA and GFP-GFP was significant (Mann-Whitney test, p-value <0.001; KS-statistic: 0.14, p-value: 2.17e-13).

In summary, both the acquisitions of endogenous and recombinant Bassoon strongly support the idea that Bassoon is oriented in a specific way at the AZ such that the N-terminus is closer to the SV while the C-terminus closer to the plasma membrane. In addition, due to constraints arising from incomplete labeling and photodamage affecting DNA strands, measurements obtained via NND tend to be overestimated. Consequently, our results indicate that the distance between the N-terminal and C-terminal regions of Bassoon is likely closer to 26 nm.

## Discussion

The cytomatrix assembled at active zones is one of the intricate elements of presynaptic nerve terminals and a hallmark of presynaptic ultrastructure. Its molecular composition and the individual functions of its constituent proteins have been extensively studied. The macromolecular complex formed by these proteins is often regarded as a scaffold or nano-machine (Schoch and Gundelfinger, 2006; Körber and Kuner, 2016; Tabares and Rizzoli, 2022). The first step towards elucidating the workings of this supramolecular complex is defining the location of its component parts relative to each other, ideally at the sub-molecular level, e.g. by locating individual domains or termini of proteins. Here, we visualize directly a spatial segregation between the N- and the C-terminus of the active zone scaffolding protein Bassoon at synapses in cultures neurons using MINFLUX.

### Visualizing the distribution of Bassoon termini with 5 nm precision in 3D

As an emerging technique, MINFLUX has been used to locate distinct types of vesicles at presynaptic terminals in cultured hippocampal neurons, revealing that ATG9-containing vesicles are distinct from synaptophysin containing synaptic vesicles, although at the confocal level these types of vesicle markers overlap entirely (Binotti et al., 2024). MINFLUX has also revealed the distribution of synaptic proteins in mouse brain tissue, including the presynaptic vesicular glutamate transporter VGluT1, the postsynaptic scaffold protein PSD95 and AMPA-receptors (Moosmayer et al., 2024). In addition, MINFLUX has revealed the distribution of presynaptic proteins in the highly specialized photoreceptor ribbon synapse terminals (Grabner et el., 2022). In contrast, MINFLUX had not been used previously to visualize and distinguish sub-molecular sites of proteins at the synapse. MINFLUX microscopy has been used to measure intramolecular distances with 1 nm precision on biochemically purified samples, where individual molecules are sparsely distributed (Sahl et al., 2024). The same study also visualized the arrangement of lamin proteins in intermediate filaments and that intramolecular distances in Lamin proteins can be directly quantified in the nuclei of transfected osteosarcoma cells. Thus, we wondered if two distinct regions of the scaffolding protein Bassoon can be resolved in the crowded environment of the active zone. Our study shows that MINFLUX indeed can achieve this. Moreover, a triple label MINFLUX analysis visualized the N-terminal region of Bassoon, the C-terminus of Bassoon as well as postsynaptic Homer and the location of the synaptic cleft simultaneously in 3D. All data sets revealed a spatial segregation of N- and C-terminal signals, with the majority of the C-terminal signals closer to the synaptic cleft than the N-terminal signals.

Our data are consistent with predictions based on electron microscopy and on STORM imaging. An early electron microscopy study detected immunogold signals for the sap7f-epitope of Bassoon (amino acids 738-1035 of rat Bassoon) between one and three synaptic vesicle diameters away from the active zone plasma membrane (Sanmarti-Vila et al., 2000). Another electron microscopy study performed on high-pressure frozen material revealed a peak of immunogold labelling for this epitope at 70 nm from the active zone membrane (Siksou et al., 2007). A third immunogold electron microscopy study revealed that C-terminal epitopes of Bassoon were found 35-37 nm away from the active zone membrane (Limbach et al., 2011). While the methods and the types of synapses differed across these studies, the results do suggest that the C-terminus of Bassoon is located closer to the plasma membrane than the N-terminus. A super-resolution light microscopy study using STORM on mouse brain sections showed that the N-terminal area of Bassoon (the epitope corresponding to amino acids 738-1035 in rat Bassoon) is – on average – farther away from postsynaptic Homer than the C-terminus of Bassoon (Dani et al., 2010). This study obtained a precision of localization of 14 nm in xy and 35 nm in z. The results were obtained by comparing sections stained for Bassoon-N and Homer with sections stained for Bassoon-C and Homer. Our MINFLUX study supports and extends the data obtained from STORM analysis, and directly visualizes the distribution of N-Bassoon and Bassoon-C within individual presynaptic terminals, as well as the distribution of N-Bassoon, Bassoon-C and Homer at individual synapses, with a precision of localization of around 5 nm in 3D.

### Strengths and limitations of our approach

We took several measures to control and validate our conclusions. First, we visualized endogenous Bassoon in naive neurons as well as full-length recombinant Bassoon in transfected neurons. Second, we employed three labelling approaches, including i) indirect labelling with primary and secondary antibodies, ii) indirect labelling with nanobodies as secondary reagents to reduce the linkage error (i.e. the error introduced by the distance between the fluorophore and the target protein), and iii) nanobodies as primary reagents directly binding to tags in recombinant Bassoon to minimize the linkage error. Third, we used both spectral demixing of fluorophores for simultaneously labelled epitopes and DNA-PAINT for successively labelled epitopes. Moreover, we performed negative-control experiments where we double labelled Bassoon using two antibodies detecting the same epitope. We performed quantitative analyses confirming that N-terminal signals and C-terminal signals represented similar numbers of localization sites, ruling out concerns over differential epitope accessibility. Finally, we implemented and verified a reliable drift-correction protocol necessary for DNA-PAINT approaches, where a washing step has to be performed between MINFLUX recordings. To verify that we exclusively imaged Bassoon at glutamatergic synapses on dendritic spines we included confocal actin staining (revealing spines) and Homer or Shank2 as molecular markers for glutamatergic spine synapses in each experiment. Together, our results indicate that this strategy was successful.

The limitations inherent in our imaging approaches are important to consider. Dual fluorophore imaging at the nanometer scale is prone to Förster Resonance Energy Transfer (FRET) between fluorophores, which can occur at distances less than 10 nm, hindering the detection of closely spaced molecules (Helmerich et al., 2022). Thus, some colocalizations may go undetected and distances may be overestimated. We addressed this known limitation, first by choosing synapses that presented adequate spectral separation, and, second, by including DNA-PAINT as a complementary approach, which circumvents FRET entirely by temporally separating the fluorophores (Ostersehlt et al, 2022). Our control experiments also revealed an important limitation to our DNA-PAINT approach: when we imaged the N-terminus of Bassoon twice in a row using DNA-PAINT and performed drift-correction we obtained an NND of 21.50 nm, despite expecting a distance near the minimum resolution distance. Since our drift correction protocol preserved localization accuracy during washing steps, additional factors may have contributed to this discrepancy. A detailed analysis of our control data revealed that, in the second acquisitions, the number of localizations was consistently lower than in the initial acquisition of the same target. While the exchangeable pool of imager strands combats the problem of photobleaching, it has been demonstrated that DNA docking strands are also susceptible to damage from reactive oxygen species (ROS), resulting in a declining number of available docking sites over time (Blumhardt et al., 2018, Soeller et al., 2025). Docking sites in close proximity to the fluorophore are most likely to experience damage, with the introduction of additional spacers to the imager shown to reduce the effect (Blumhardt et al., 2018, Clowsley et al., 2021). This is of particular relevance to experiments where the same docking strands are imaged repeatedly, as in the control measurements described here. While experiments focusing on the two terminals of Bassoon do not show a consistent count reduction in the second measurement, the second measurements in control experiments largely show fewer localizations than the first, and do not show perfect correlation in binding sites which might account for the artifactually large NND calculated for the synaptic controls.

Thus, our study has allowed us to validate and refine our understanding of Bassoon’s spatial organization within the active zone. In addition, it also underscores the current technical constraints associated with the two strategies employed to achieve nanometer-scale resolution in three-dimensional imaging. The primary limitation of spectral demixing, as described by Helmerich et al. (2022), lies in the potential interactions between fluorophores, which can impact localization accuracy. Similarly, exchange-PAINT is constrained by the susceptibility of DNA strands to photodamage. In both cases, these effects can lead to a loss of localizations, artificially increasing the measured distance between two populations while simultaneously reducing localization density. In our study, we used Gaussian Mixture Model analysis to describe the distribution of nearest neighbor distances. We assume that the first peak in the histogram represents the closest distance observed between two Bassoon signals, yielding a reliable estimate of the nearest neighbour distance even if localizations are lost. However, our estimate of the number of Bassoon copies is inevitably an underestimate of the actual number of copies. An alternative approach to overcoming these limitations and maximizing the potential given by MINFLUX microscopy in order to address biological questions that cannot be resolved using techniques such as STORM and PALM would be to combine spectral demixing with DNA-PAINT. This strategy was proposed by Gimber et al. (2022) under the name SD-DNA-PAINT and the authors achieved to image simultaneously 3 different targets using DNA-PAINT and spectral demixing on a d-STORM setup. Applying this strategy to MINFLUX microscopy would accelerate acquisition time, reducing the likelihood of damaging the DNA strand responsible for nanobody binding. In addition, the elimination of liquid exchange would eliminate potential drift between sequential acquisitions. This will require the development of new probes and fluorophores in the near future.

### Quantitative insights

Using DNA-PAINT and 3D MINFLUX we detected on average 155 copies of endogenous Bassoon at an active zone. This is, to our knowledge, the first estimation of the number of Bassoon copies at the active zone using SMLM imaging techniques. This number is lower than predicted based on biochemical experiments, which suggested that 450 copies of Bassoon reside – on average – at a synapse (Wilhelm et al., 2014). As discussed above, our estimate is probably an underestimate of the actual number of Bassoon copies. Of note, we selectively determined this number for a subset of synapses, i.e. glutamatergic presynaptic terminals contacting dendritic spines.

The unprecedented precision of localization of 3D-Minflux unveiled the spatial arrangement of Bassoon molecules in the active zone. The measured NND between two N-terminal Bassoon regions consistently showed a preferred interspacing between two N-termini of ∼10 nm, with a median of 14.60 nm. Notably, this spacing was identical when comparing two C-termini, suggesting equivalent arrangement and constraints on both termini. These results suggest that one site containing Bassoon is generally present within 10 nm of the next. Again, we found similar intermolecular distances for the recombinant protein with a median of the distance between two GFP-tags of 16.10 nm and between two ALFA-tags of 15.10 nm. In addition, both for the endogenous proteins and for the recombinant protein the N-terminal area occupied a similar volume as the C-terminal area. While this does not allow us to make any inference regarding the shape of individual Bassoon molecules it does indicate that the N-termini and the C-termini have virtually the same spacing between neighboring Bassoon molecules and argues for an orderly arrangement of the protein. This is unlike, for example, the arrangement of the filamentous active zone protein Bruchpilot at the Drosophila neuromuscular junction (Fouquet et al., 2009). At these presynaptic terminals the N-terminus of Bruchpilot is oriented towards the plasma membrane while the C-terminus of is oriented towards synaptic vesicles. This vertical orientation at the active zone is reminiscent of Bassoon, albeit with opposite directions of the termini. Of note, the membrane proximal terminus of Bruchpilot occupies a smaller area than the SV-proximal C-terminus, representing the shape of the Drosophila T-bar. Our data argue against such an arrangement of Bassoon because both termini of Bassoon cover similar volumes and areas.

Imaging the N-terminus twice revealed a 10 nm periodicity in the distribution of the separation distances, which we interpret to reflect the distance of the first, second, third and fourth closest Bassoon. In contrast, the distribution of separation distances between the closest N- and C-terminal regions of Bassoon fits more closely to a Gaussian profile with a mean at around 26 nm, with the interesting exception of an additional peak at 14 nm. Although it is not possible to definitively assign a specific N-terminal to its corresponding C-terminal within the same Bassoon molecule, this pattern may point to a diversity of conformations—some Bassoon either more folded or showing more intertwined 3D configurations between them—within specific regions of the AZ.

In their study, Dani et al revealed that Bassoon-C was, on average, located 30 nm closer to Homer than N-Bassoon, suggesting a specific orientation of Bassoon at the active zone (Dani et al, 2010). To obtain these results they co-stained Homer and the N-terminal area of Bassoon as reference points and determined the center of the synaptic cleft as the midpoint between N-Bassoon and Homer-C. Their study represented the first attempt to characterize the spatial organization of presynaptic active zone proteins at a resolution approximately tenfold greater than the resolution of conventional light microscopy. Here, we visualize the arrangement of the N-terminal and C-terminal of Bassoon at the AZ with 5 nm 3D localization precision. Of note, while our approach can visualize the distinction between the two regions of Bassoon directly, and visualize the distribution of these regions at the active zone on a single molecule basis, we cannot assign one certain N-terminal signal to its corresponding C-terminus in the same molecule.

Interestingly, the NND for the N-terminal versus C-terminal area of the endogenous protein (28.80 nm) was slightly larger than the NND for recombinant protein, which was 25.90 nm. For endogenous Bassoon, a combination of primary antibodies and secondary nanobodies was used, which may result in artificial shrinkage or growth of the spatial arrangement. Conversely, for GFP-Bassoon-ALFA, nanobodies against GFP and ALFA were used in order to minimize linkage error. This could explain the shorter distance obtained for GFP-Bassoon-ALFA than for endogenous Bassoon. However, considering limitations such as labelling efficiency and photodamage, our results suggest that the actual distance between the N-terminus and the C-terminus of the Bassoon is shorter than previously estimated and cannot exceed 26 nm. All three predicted coiled-coil domains of Bassoon are located downstream of amino acid 738 (tomDieck et al., 1998). These domains are predicted to be elongated structures (Gundelfinger et al., 2016). Modeling by Alphafold 2 (Jumper et al., 2021) has predicted the longest of these three coiled-coil domains to be 17 nm. Thus, the major determinants for the length of Bassoon appear to be located downstream of amino acid 738. Moreover, the N-terminal area upstream of amino acid 738 may be flexible and fold back towards the C-terminus.

Together, by employing and optimizing novel microscopy and labeling techniques, our study is setting a new standard for dissecting the complex molecular organization of excitatory synapses, which remains an important research subject for molecular and cellular neuroscience.

## Methods

### Plasmid construct

The dually tagged full-length rat Bassoon construct was created in the ampicillin-resistant pCS2^+^ vector backbone and designed to include both an intramolecular mEGFP tag and the ALFA tag (PSRLEEELRRRLTE; Götzke et al., 2019) fused to the C-terminus of Bassoon. The intramolecular tag was created by gene synthesis in a way that its coding sequence, after insertion into the HindIII site of rat Bassoon, was preceded by amino acids 1-97 of Bassoon and was followed by amino acids 95-3938 of Bassoon (Ghelani et al., 2021).

### Primary hippocampal neuron cultures and transfection

Primary hippocampal neurons were prepared in Kaech and Banker’s mode on an astrocyte feeder layer (Kaech and Banker, 2007). The astrocytes were isolated from cortices of E19 Wistar rat in ice cold Hank’s Balanced Salt Solution (HBBS, Gibco) supplemented with 100 U/ml penicillin/streptomycin (Gibco) and dissociated by trypsination with 0.25% Trypsin (Gibco). The cells were cultured two weeks in T75 Flasks Minimum Essential Media (MEM, Gibco) supplemented with 10% (v:v) Horse serum (Gibco), 2 mM of L-Glutamine (Gibco) and 4.65 g/L of D-(+)-glucose (Sigma). After two weeks the astrocytes were harvested and plated at a density of 100 000 cells per 60 mm Petri dishes coated with 0.1 mg/ml of Poly-L-lysine (Sigma). The hippocampal cells were plated on glass coverslips with paraffin dots coated with 1% (v:v) polyethyleneimine (PEI, Sigma) in Dubelcco’s Modified Eagles Medium (DMEM, Gibco) supplemented with 10% (v:v) of fetal bovine serum (FBS, Gibco), 2 mM of L-Glutamine (Gibco) and 100 U/mL penicillin-streptomycin at a low density of 10 000 cells per cm^2^. After 2h of incubation at 37°C and 5%CO_2_ and 2 coverslips were flipped into the dish containing the astrocyte feeder layer. The neurons were maintained in Neurobasal medium (Gibco) supplemented with 2% B27 (Gibco) and 2 mM L-glutamine. After 3 days *in vitro* the cultures were treated with 4.9 µM of cytosine α-D-arabinofuranoside hydrochloride (AraC, Sigma) to stop the proliferation of glial cells.

Primary hippocampal neurons were transfected at 2 days *in vitro* with GFP-Bassoon-ALFA plasmid using Calcium phosphate transfection method. Briefly, 2 mL of pre-warmed maintenance media was added to the 60 mm Petri dish before to start the transfection. For each 25 mm coverslip 3 µg of DNA were directly mixed with 2M CaCl_2_. The solution was then diluted with water to reach a final concentration of 250 mM CaCl_2_. The transfection buffer (274 mM NaCl, 10 mM KCl, 1.34 mM Na2HPO4, 15 mM D-glucose, 42 mM HEPES, pH 7.09) was added dropwise while vortexing to the solution in a final dilution 1:1. The mix was incubated 20 min at room temperature in dark. 900 µL of the preconditioned medium from the dish was transferred in a well of a 6 well plate and the coverslip was flipped neurons on the top on the well. 100 µL of the mix solution DNA/CaCl2 and transfection buffer was added drop by drop on the well and the cells were incubated 1 h at 37°C and 5% CO2. After checking the formation of a precipitate on the cells, the well was washed 2 times with prewarmed HBSS with osmolality adjusted to the osmolality of the maintenance medium to wash the precipitate. The cells were incubated 10 min at 37°C and 5%CO2 in the HBSS and then the coverslip was flipped back to the original Petri dish.

### Immunostaining of hippocampal neurons

Hippocampal neurons were fixed at day in vitro 14 in 2.4% paraformaldehyde in phosphate buffered saline (PBS) for 30 min at room temperature. The cells were permeabilized 5 min with 0.4% Triton (Sigma). The cells were rinsed in PBS and the aldehyde fluorescence was quenched with 50 mM NH4Cl for 10 min and then the cells were blocked 45 min in 2% Bovine Serum Albumin (Sigma). The cells were then incubated overnight at 4°C with primary antibodies diluted as follows: monoclonal rabbit anti Homer1 (1:1000, Synaptic Systems, #160 008, RRID:AB_2832230), monoclonal mouse anti Bassoon raised against amino acids 738-1035 of rat Bassoon (1:1000, Enzo, SAP7F407, RRID:AB_422275), monoclonal rabbit anti Bassoon raised against amino acids 756-1000 of rat Bassoon (1:1000, Synaptic Systems, #141 118, RRID:AB_2864773), polyclonal rabbit anti Bassoon raised against the carboxy terminus of Bassoon (1:1000, Synaptic Systems, #141 003, RRID:AB_887697), polyclonal guinea-pig anti Shank2, Synaptic Systems, #162 204, RRID:AB_2619861), guinea pig anti VGluT1 (1:1000, Synaptic Systems, #135 304, RRID: AB_887878) After washing in PBS the samples were incubated 45 min at room temperature with secondary antibodies: anti rabbit coupled to Cy3 (1:1000, Jackson ImmunoResearch Labs, 711 165 152, RRID:AB_2307443),anti guinea-pig coupled to Cy3 (1:1000, Jackson ImmunoResearch Labs, 706 165 148, RRID:AB_2340460), phalloidin-AF488 (1:400, Invitrogen, A12379).

For spectral demixing MINFLUX the cells were stained with goat-anti-ms-Flux-680 (Abberior, 1:200) and goat-anti-Rb-Flux-640 (Abberior, 1:200) in 2% BSA in same times as the other stainings. For DNA-PAINT the cells were blocked 30 min in antibody incubation buffer (Massive Photonics) and then incubated 1h with nanobodies in antibody incubation buffer: Nanobody against mouse with docking site 1 (1:100, MASSIVE-SDAB 2-PLEX, Massive Photonics), nanobody against rabbit with docking site 2 (1:100, Massive Photonics MASSIVE-SDAB 2-PLEX, Massive Photonics) and nanobody against ALFA tag and coupled to docking site 3 (1:200, MASSIVE-TAG-Q ANTI-ALFA, Massive Photonics), nanobody against GFP and coupled to docking site 4 (1:100, MASSIVE-TAG-Q ANTI-GFP, Massive Photonics).

### U2OS cell culture and preparation

U2OS-CRISPR-NUP96-mEGFP cells (Thevathasan er al., 2019, Cytion #300174) were cultured in McCoys 5A medium (modified) (with 3.0 g/L Glucose, stable Glutamine, 2.0 mM Sodium pyruvate, 2.2 g/L NaHCO3, Cytion, #820200a). Cells were seeded on #1.5 coverslips to reach 40% confluence on the day of fixation.

### U2OS cell fixation and staining

Cells were fixed with 2.4% PFA for 30 minutes followed by a 3-minute extraction with 0.4% Triton X-100. Samples were quenched with () NH4Cl before blocking with antibody incubation buffer (Massive Photonics) for 30 minutes. The anti-GFP-X2 nanobody (1:200, MASSIVE-TAG-X2 anti-GFP, Massive Photonics) was made up in antibody incubation buffer and incubated at room temperature for 1 hour. The sample was then washed 3 times with PBS before further preparation for MINFLUX imaging.

### Gold beads treatment for active sample-stabilization for MINFLUX microscopy

After the samples were washed in washing buffer (Massive Photonics) the cells were incubated 5 min with 150 nm colloidal gold beads (BBI Solutions, EM. GC150) and the excess of gold beads was washed away with three PBS rinses. The gold beads were then stabilized by incubation with 10 mM MgCl_2_ for 5 min, which was then replaced with 1 mg/ml of Poly-L-lysine (Sigma) for 5 min.

For spectral demixing MINFLUX, the coverslips were mounted on a microscopy slide in a STORM blinking buffer (pH 8.0) composed of 50 mM Tris-HCl, 10 mM NaCl, 10% Glucose (w:v), 64 µg/ml catalase (Sigma), 0,4 mg/ml glucose oxidase (Sigma) and 53-63 mM MEA (Sigma). For DNA-PAINT, samples were mounted in a magnetic imaging chamber in which the cells were incubated with the complementary DNA imager strand coupled to the fluorophore ATTO 655. The imager strand was made up in imaging buffer (Massive Photonics) to concentrations between 500 pM and 2 nM.

### MINFLUX 3D microscopy

Samples were imaged on two Abberior Instruments MINFLUX microscopes. Details of the system layout and components are described in Schmidt et al., 2021. Briefly, the microscopes were built on motorized inverted microscope bodies (IX83, Olympus) with a CoolLED pE-4000 (CoolLED) for epifluorescence illumination, and equipped with 405 nm, 485 nm, 561 nm, and 642 nm laser lines, as well as a 980 nm IR laser and xyz piezo (Piezoconcept) for the active sample stabilisation system. The reflection from the 980 nm laser were recorded by the dedicated stabilisation system and used to assess sub nanometer shifts in the sample position. These deviations were corrected actively during the measurement with the xyz piezo stage. Both systems contain a 642 nm MINFLUX laser. Images were acquired with a 100x 1.45 NA UPLXAPO oil objective (Olympus) or 60x 1.42 NA UPLXAPO oil objective (Olympus). The synapses were selected in the confocal mode based on the fluorescence of the GFP and the post synaptic protein Homer 1 for the recombinant Bassoon, or the actin staining and the co-staining of the post synaptic proteins Homer 1 or Shank2 for endogenous Bassoon. The three-dimensional spectral demixing MINFLUX images (3D-SD MINFLUX) were recorded with a pinhole diameter of 0.77 Airy Units, a MINFLUX laser power of 5-6 % and a spectral window from 682 nm – 780 nm for detector 1 and a spectral window from 651 nm – 690 nm for detector 2.

The three-dimensional (3D) DNA-PAINT MINFLUX images of Bassoon arrangement were recorded with a pinhole diameter of 0.6 Airy Units, a MINFLUX laser power of 16 % and a spectral window of 654 nm to 770 nm.

For MINFLUX beamline monitoring (mbm), 7-9 gold beads were selected as reference markers and used to correct any system drift over the course of the measurement.

After acquisition of the first target, the samples were washed one time in 1X washing buffer (Massive Photonics) and then incubated with the second imager strand.

For both SD MINFLUX and DNA-PAINT MINFLUX, imaging was performed with the default 3D imaging sequence supplied with the system. In brief, the sequence defines parameters used in the iterative localising process of MINFLUX, culminating in a final pair of lateral and axial iterations with a TCP (targeted coordinate pattern) diameter of 40 nm, and minimum photon thresholds of 100 and 50 photons, respectively. Key parameters are given in the table below.

| Iteration | L (nm) | TCP | Photon Threshold | cfr check | Dwell time (s) | Pattern Repeats | Background threshold (Hz) | Power Factor |
| --- | --- | --- | --- | --- | --- | --- | --- | --- |
| 0 | 288 | Hexagon | 160 | - | 0.001 | 1 | 15000 | 1 |
| 1 | 288 | z-line | 400 | - | 0.001 | 1 | 15000 | 1 |
| 2 | 288 | Square | 100 | - | 0.001 | 5 | 10000 | 1 |
| 3 | 288 | z-line2 | 50 | - | 0.001 | 5 | 10000 | 1 |
| 4 | 151 | Square | 67 | 0.9 | 0.001 | 5 | 10000 | 2 |
| 5 | 151 | z-line2 | 33 | - | 0.001 | 5 | 10000 | 2 |
| 6 | 76 | Square | 67 | 0.8 | 0.001 | 5 | 10000 | 4 |
| 7 | 76 | z-line2 | 33 | - | 0.001 | 5 | 10000 | 4 |
| 8 | 40 | Square | 100 | - | 0.001 | 5 | 10000 | 6 |
| 9 | 40 | z-line2 | 50 | - | 0.001 | 5 | 10000 | 6 |

### Spectral demixing MINFLUX analysis

Spectral unmixing was performed on pyMINFLUX. The dcr is computed as a weighted average using the effective counts periphery (eco) applied to the initial iterations. A Gaussian mixture model (GMM) with two components is fitted to the result. Based on the parameters obtained from the GMM fitting, a color assignment is performed. All fluorophores belonging to the same trace are assigned a specific color, determined by the predominant classification within that trace. Figures with each trace were made using Paraview 5.8.1. Figures with the centroids localization of each burst were generated with PyMINFLUX and PYMEVisualize.

### 3D DNA-PAINT MINFLUX analysis

Fiducials positions used for MINFLUX beamline monitoring (MBM) during acquisition were used to calculate a 3D rigid transform to map the post-wash data set to the pre-wash data, correcting for any nanometer scale shifts introduced during buffer exchange. Quality of the selected beads was assessed by inspecting the bead trajectories and localisation precisions over the acquisition time. The average position over the first 5 localisations for each bead was calculated and used as the fiducial position for the calculation of the transformation. Translation and rotation were calculated using linear algebra and applied to the already MBM corrected post-wash dataset, resetting position 0 for the beads to match one another, with the varying drift during acquisition still accounted for with the routine MBM correction.

The mean absolute error of bead localization after drift correction was calculated as:

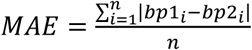

with *bp*1_*i*_ and *bp*2_*i*_ corresponding to the position of the bead at the start of the first and second measurements respectively.

To compensate for the refractive index mismatch, data was scaled in z by a factor of 0.7, as discussed in Gwosch et al. (2020) trace IDs containing fewer than 3 localizations were filtered out prior to calculation of the centroids of each localisation burst. Centroids were merged into units as described in Gürth et al. (2024) with a conservative value of 6.5 nm chosen as the merge distance, calculated using the mean raw localisation precision in each axis plus 1 standard deviation, as follows: ∼√(2σx+ 2σy+ 2σz). Synapses were identified using DBSCAN with an epsilon value of 100 nm and minimum point count of 20. Nearest neighbor distances were then calculated within and between populations using a kd tree algorithm. For the calculation of nearest neighbors, the convex hull of the point cloud after filtering and synapse identification was calculated and the volumes between the two populations compared. In order to avoid artificial over estimation of NND distributions, the synapse with the smaller convex hull volume was used as the test points, and the larger the points to be searched. This reduces the impact of any issues in the cluster identification or mismatches in region quality.

The Kolmogorov-Smirnov test was used to compare the distributions of nearest neighbor distances between GFP-AFLPA vs GFP-GFP and between N-N vs N-C terminals to determine if they were drawn from the same distribution. The test statistic (D) was 0.14, with a corresponding p-value of 2.17e-13 when comparing GFP-ALFA vs GFP-GFP distances, and D = 0.13, p-value of 8.36e-17 when comparing N-N vs N-C distances, respectively. Given a significance level of α = 0.05, there is a significant difference between the distributions in both cases: GFP-AFLPA vs GFP-GFP and N-N vs N-C.

Number of Bassoon copies per active zone were calculated using PyMINFLUX (Ponti et al, 2023) and python. As for DBSCAN analyses, every centroid of burst located within a distance below the raw localization precision were counted as a unit reflecting the same Bassoon copy. Figures of centroids of each localization burst were generated with PyMINFLUX and PYMEVisualize (Marin et al, 2021). The volume of expression of each target was quantified using Point Cloud Analyst (PoCa) software (Levet et al, 2015; Levet et al, 2019).

### Gaussian Mixture Modelling

We employed a Gaussian Mixture Model (GMM) framework to model the underlying distribution of the data and to infer the optimal number of components. Model estimation was performed using the Expectation-Maximization (EM) algorithm as implemented in the scikit-learn library. To select the optimal number of mixture components (K), we fit GMMs across a range of candidate models (K=1 to 20) using full covariance matrices. Model quality was evaluated using Bayesian Information Criterion (BIC). The model with the lowest BIC was selected as the optimal fit. To assess the stability and robustness of the selected model, we employed a bootstrap resampling approach. Specifically, we generated 100 bootstrap samples from the original dataset by sampling with replacement. For each bootstrap replicate, we refit GMMs across the same candidate range and recorded the value of K that minimized the BIC. The most frequent value of K was selected as the best candidate. All analyses were performed in a custom code written in Python (version 3.11) using NumPy and scikit-learn.

### Quantification of Bassoon overexpression

Quantification of Bassoon intensity in epifluorescence data was done using ImageJ and GraphPad. Images were taking on a widefield Zeiss Axio Imager.Z2 epifluorescence microscope equipped with the 16-bit ORCA-flash4.0 V2 digital CMOS camera (Hamamatsu) and ZEN Blue software version 2.3. Bassoon fluorescence intensity in synapses expressing recombinant and synapses expressing only endogenous Bassoon was determined using a mask between Bassoon and VGluT1 and a mask between GFP and VGluT1 to select presynaptic Bassoon and to exclude any potential ectopic expression of recombinant Bassoon. The GFP:VGluT1 mask was then applied to synapses selected with Bassoon:VGlUT1 mask and Bassoon intensity was measured in synapses within the GFP:VGluT1 mask (expressing recombinant and endogenous Bassoon) and outside the mask (expressing endogenous only). The quantification was done on N = 3 independent cultures preparation and for each cultures n = 30 cells. To test for normality of the data we applied Shapiro-Wilk test.

## Acknowledgements

We thank Tanvir Rahman Shaikh, Senior Research Engineer at the Ulmeå Centre for Electron Microscopy, Ulmeå University for his help in analyzing 3D data, in particular writing a python script to determine the number of Bassoon copies per active zone. We thank Nina Dankenbrink-Werder and Lisa-Marie Hartmund for excellent technical assistance.

