## supplemental data for "3D MINFLUX combined with DNA-PAINT resolves the arrangement of Bassoon at active zones"

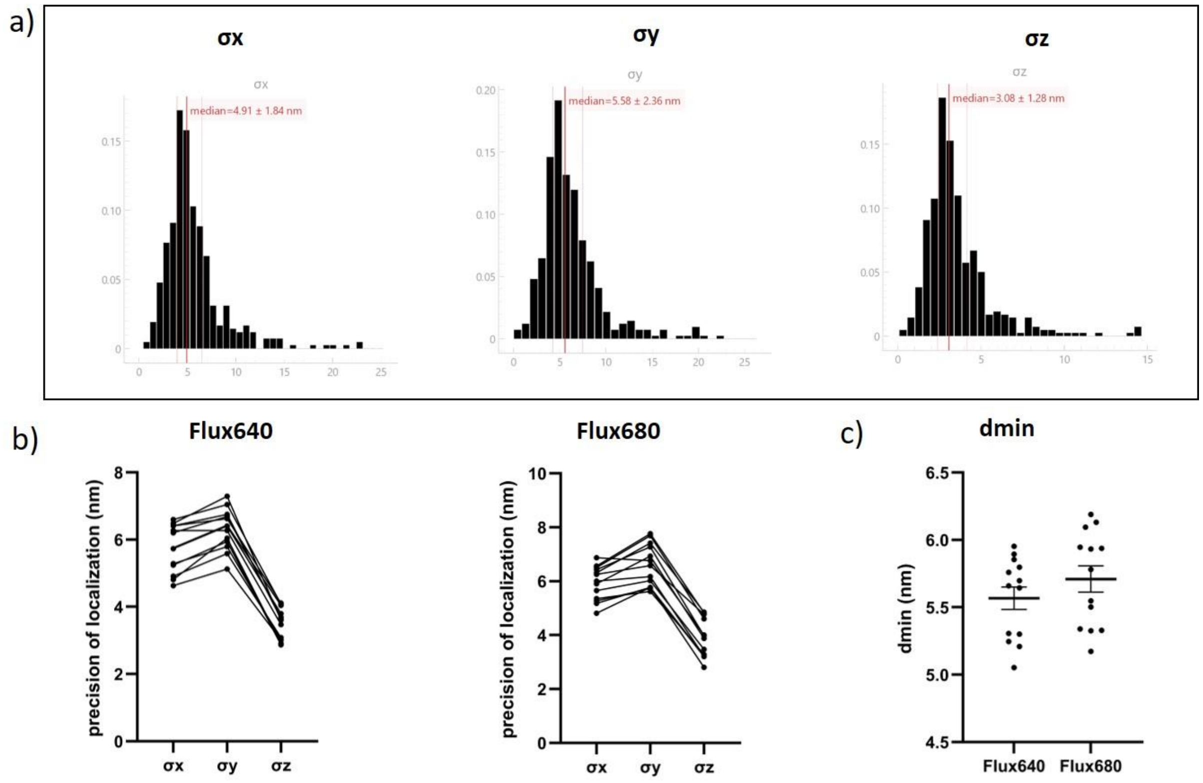


**Figure S1: precision of localization and resolution of the Flux640 and Flux680 in 3D Minflux data.** a) representative example of precision of localization x, y, z in one dataset of N-Bassoon with Flux640. b) precision of localization for each datasets and imagers. c) dmin calculated as $=\sqrt{(2\sigma x+2\sigma y+2\sigma z)}$ for Flux640 (5.57 ± 0.08 nm; mean ± SEM) and Flux680 (5.71 ± 0.10 nm; mean ± SEM)

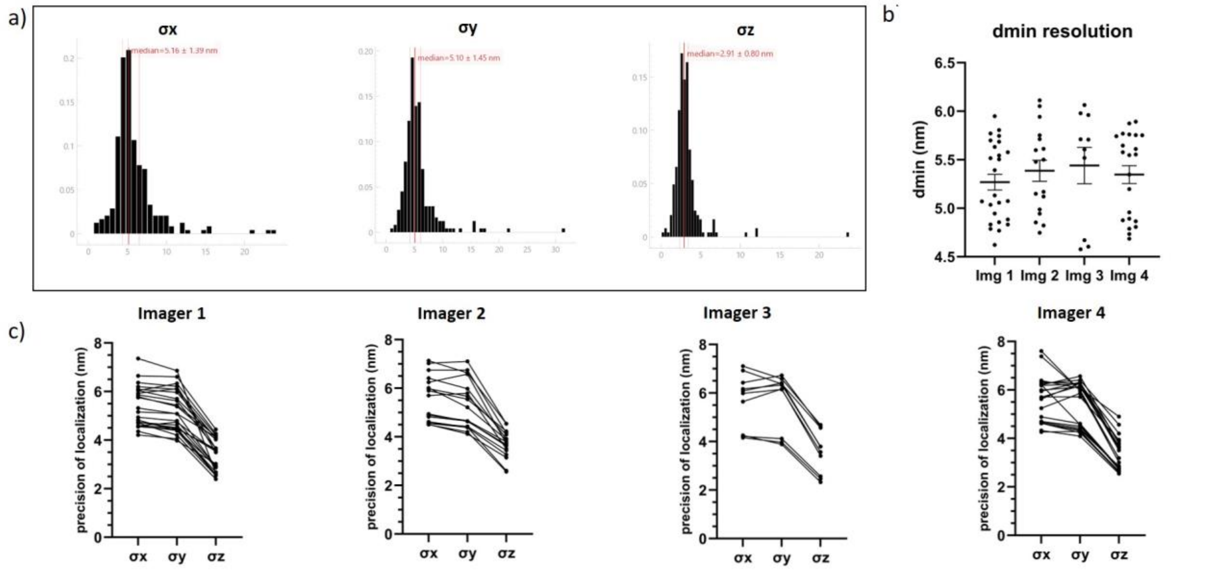


**Figure S2: precision of localization and resolution of the DNA-PAINT 3D Minflux data.** a) representative example of precision of localization in x, y, z in one dataset of N-Bassoon using Imager 1. b) dmin calculated for each datasets using Imager 1 (Img 1), 2, 3 or 4 as dmin $=\sqrt{(2\sigma x+2\sigma y+2\sigma z)}$ . c) precision of localization for each datasets and imagers

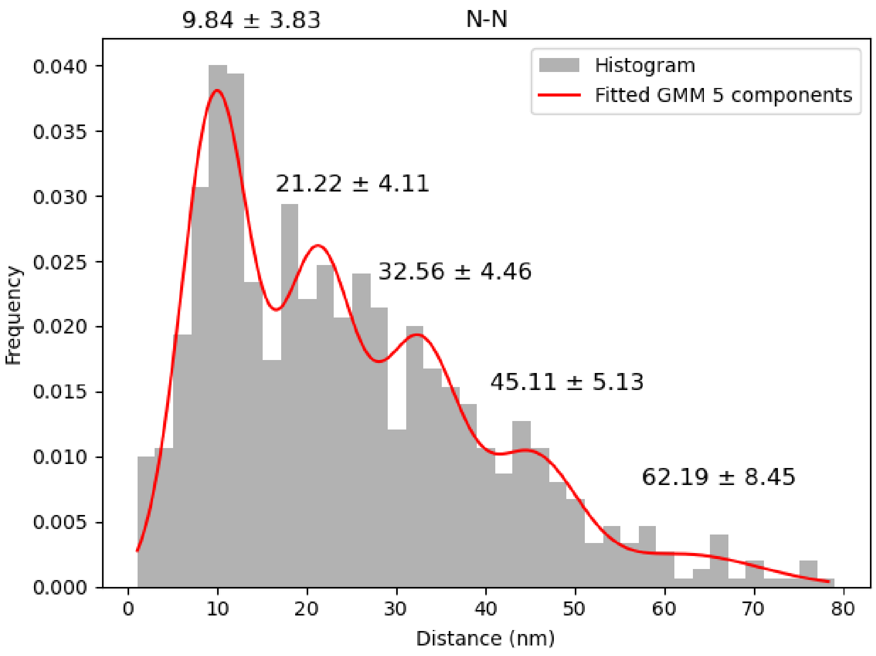


**Figure S3: Analysis of the distribution of nearest-neighbour distances between twice-imaged N-termini of Bassoon using a Gaussian Mixture Model.** A periodicity of 11 nm is found between each peaks.

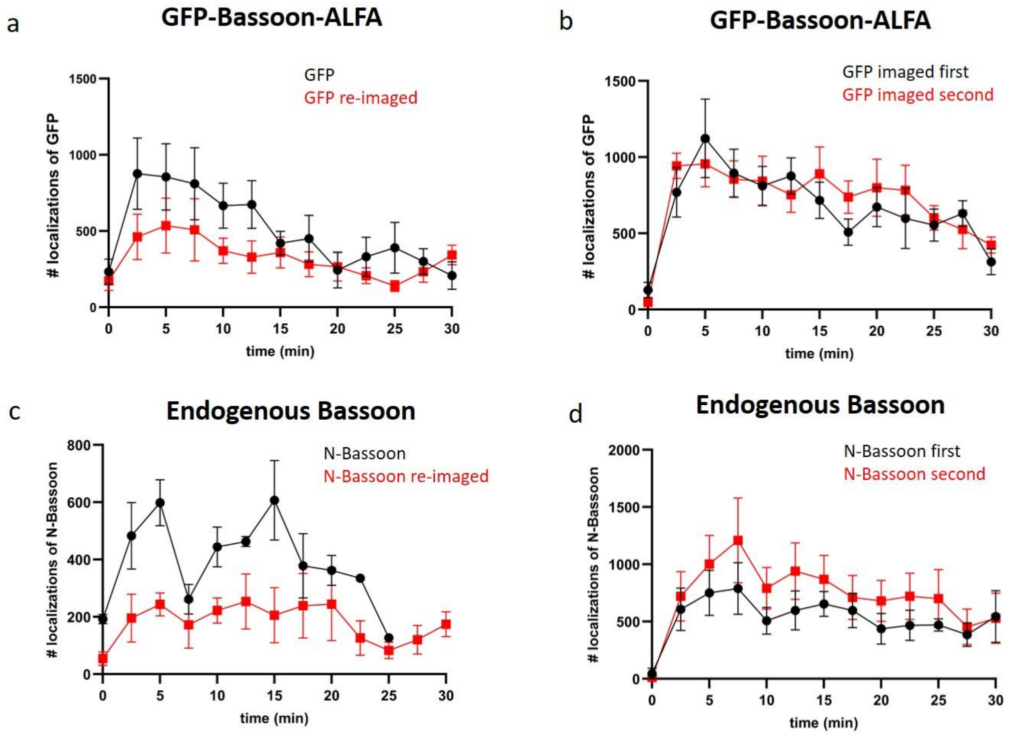


**Figure S4: Monitoring the variation in loc count over time may indicate signs of photodamage.** a) Temporal variation in GFP localizations during the first acquisition (black) and subsequent recording (red). b) Temporal variation in GFP localizations in GFP-Bassoon-ALFA, where GFP was acquired either first (black) or after ALFA (red). c) Temporal variation in N-Bassoon localizations across the first acquisition (black) and second recording (red). d) Temporal variation in N-Bassoon localizations in N-Bassoon/Bassoon-C acquisitions, with N-Bassoon recorded either first (black) or after Bassoon-C (red).

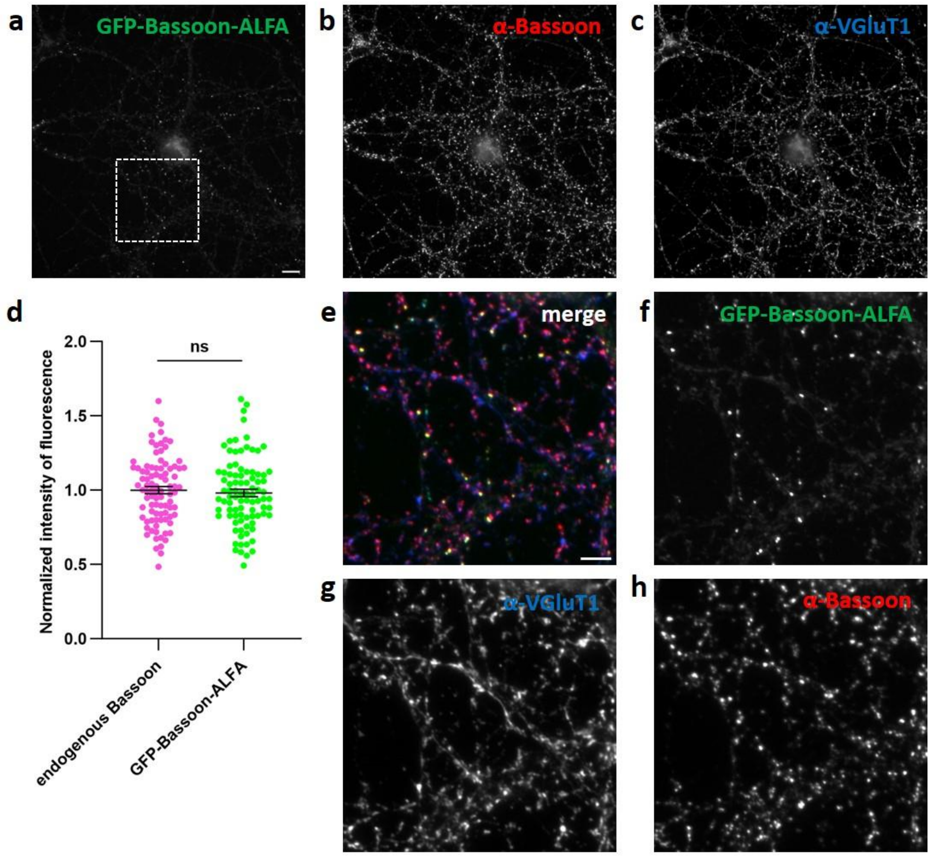


**Figure S5: Expression of recombinant Bassoon shows no increase of Bassoon expression at the synapse.** Epifluorescence microscopy of rat hippocampal neurons at DIV14 transfected with GFP-Bassoon-ALFA (a) and co-stained with anti-N-Bassoon (b) and anti-VGluT1 (c). d) Quantification of fluorescence from anti-Bassoon in presynaptic terminal expressing endogenous Bassoon only or GFP-Bassoon-ALFA and endogenous Bassoon showing no significant difference (mean ± SEM). N= 3 independant cultures, n = 30 images per culture. e-h) zoom of the region showed in a).
